# EVL and MIM/MTSS1 regulate actin cytoskeletal remodeling to promote dendritic filopodia in developing neurons

**DOI:** 10.1101/2021.05.26.444586

**Authors:** SS Parker, KT Ly, AD Grant, A Wang, JD Parker, MR Roman, M Padi, CW Wolgemuth, PR Langlais, G Mouneimne

## Abstract

Dendritic spines are the postsynaptic compartment of a functional neuronal synapse, and are critical for synaptic connectivity and plasticity. The developmental precursor to dendritic spines, dendritic filopodia, are highly motile protrusions that facilitate synapse formation by sampling the environment for suitable axon partners during development and learning. Despite the significance of the actin cytoskeleton in driving these protrusions, the actin remodeling factors involved in this process are not fully characterized. In this work, we identify a critical function for the Ena/VASP protein EVL in the regulation of dendritic filopodia. Amongst the Ena/VASP proteins, EVL is uniquely required for the characteristic morphology and dynamics of dendritic filopodia. Using a combination of genetic and optogenetic manipulations, we demonstrate that EVL promotes protrusive motility through membrane-direct actin polymerization at dendritic filopodia tips. EVL forms a complex at nascent protrusions and dendritic filopodia tips with MIM/MTSS1, an I-BAR protein recently discovered to be important for initiation of dendritic filopodia. We propose a model in which EVL cooperates with MIM to elongate and coalesce branched actin filaments, establishing the dynamic lamellipodia-like architecture of dendritic filopodia in developing neurons.

## INTRODUCTION

The neuronal synapse is the communication interface between neurons. Aberrant synaptic structure and connectivity is implicated in neurodevelopmental disorders, including intellectual disability, schizophrenia, and autism spectrum disorder (ASD) (Fromer et al., 2014; Gilman et al., 2011; Rubeis et al., 2014). Dendritic filopodia (DF) are actin-rich synaptic precursors that provide opportunities for new synaptic connections, and are abundant during neonatal neurodevelopment and activity-dependent plasticity (Portera-Cailliau et al., 2003; Ziv and Smith, 1996; Zuo et al., 2005). During synaptogenesis, these highly dynamic protrusions emanate from the dendritic arbor, sampling axon partners with which to establish new connections. If the axo-dendritic pairing is favored, DF are stabilized and can mature into dendritic spines, the postsynaptic compartment of excitatory synapses. Recent works suggest that the protrusion dynamics of DF influence nascent synapse formation and the capability to remodel into a spine (Carlson et al., 2011; Kayser et al., 2008; Sanchez-Arias et al., 2020). Importantly, across several neurodevelopmental conditions, an exuberance of DF, morphologically immature synapses, and altered actin dynamics is observed in patients, as well as in mouse and *in vitro* models of Fragile X syndrome, ASD, and schizophrenia (Cruz-Martín et al., 2010; Griesi-Oliveira et al., 2018; Isshiki et al., 2014; Jia et al., 2014; Sudarov et al., 2013). These conditions are frequently associated with mutations and variants in actin-associated proteins, suggesting a convergent etiological mechanism of dysregulation in actin dynamics, and highlighting the necessity of exquisitely tight control of the actin cytoskeleton for appropriate neural connectivity (Fromer et al., 2014; Gilman et al., 2011; Yan et al., 2016). As such, uncovering the actin regulators involved in the initiation and dynamics of DF informs not only the molecular basis of neuroplasticity in development and learning, but furthers our understanding of the pathophysiology of neurodevelopmental disorders.

The organization of actin in DF is defined by Arp2/3-mediated branched actin and actin filaments of mixed polarity; this cytoskeletal architecture is distinct from the parallel bundles of linear actin observed in conventional cell filopodia (Hotulainen et al., 2009; Korobova and Svitkina, 2010). Extensive works have established that Arp2/3 is required for the initiation of DF and spine morphogenesis, and that its loss or dysregulation is associated with behavioral deficits in mice (Hotulainen et al., 2009; Kim et al., 2013; Spence et al., 2016). Though necessary, Arp2/3 activity alone is not sufficient to create actin networks. Rather, Arp2/3 nucleates actin branching from pre-existing filaments, creating free barbed ends for polymerization by actin elongation factors (AEFs) (Chesarone and Goode, 2009). Two families of AEFs are found in mammalian cells – formins and Ena/VASP, which associate processively with the barbed ends of actin filaments and facilitate the addition of profilin:G-actin complexes (Chesarone and Goode, 2009). Although critically important for actin remodeling, the specific AEFs contributing to DF dynamics are not fully characterized. Notably, despite their essential functions in regulating neuronal morphogenesis and axonal growth cone filopodia (Dent et al., 2007; Kwiatkowski et al., 2007; Lebrand et al., 2004; Menon et al., 2015), and in contrast to the described role for formins in DF (Galbraith et al., 2018; Hotulainen et al., 2009; Spence et al., 2016), whether the Ena/VASP family plays a role in DF dynamics is largely unknown.

In this study, we determined that EVL is the dominant Ena/VASP paralog expressed during early synaptogenesis in cortical neurons, and is required for DF morphogenesis and protrusive motility. EVL enriches to the tips of DF in an EVH1-dependent manner, and enhances their dynamics by promoting actin polymerization. Loss of EVL by genetic knockout results in a failure of DF elongation, leaving small lamellipodia-like dendritic protrusions that are formin- and Arp2/3-dependent. Further, by acutely localizing EVL through optogenetic approaches, we demonstrated that EVL is both necessary and sufficient for DF motility. Using an unbiased proteomics approach, we identified a complex between EVL and the inverse bar (I-BAR) protein MIM/MTSS1, which interact at nascent protrusions and DF tips to promote DF initiation and motility, respectively. Together, our findings support a model where collaboration of MIM, Arp2/3, and formins at the initial protrusion provides a “hot spot” of dynamic actin, which is coalesced and elongated by the dynamic enrichment of EVL to protrusion tips, giving rise to a canonical DF.

## RESULTS

To investigate actin remodeling in DF during early synaptogenesis, we employed live-cell imaging and a quantitative pipeline to capture and define DF dynamics. DF engage in many dynamic behaviors, including initiation, protrusion and retraction. We disentangle these behaviors by extracting several metrics from tip tracking data. Absolute tip displacement in time represents the net dynamics arising from all behaviors. We established a “substantiative motility” threshold of 0.0128µm/s (one pixel displacement per 5s interval), in order to define DF or durations of time as motile or non-motile. Protrusion and retraction events, indicative of actin remodeling, are captured by the change in DF length between successive timepoints; rate is derived from the median of all instantaneous changes in length exceeding ±0.0128µm/s. Primary cortical neuronal cultures were imaged at *in vitro* day 11 (D11), which immediately precedes a developmental period of robust synaptogenesis (Fig. S1A). Due to the rapid motility of DF, we used Total Internal Reflection Fluorescence Microscopy (TIRFM) to maximize acquisition rate while minimizing phototoxicity.

### EVL is the predominant Ena/VASP paralog regulating dendritic filopodia

To determine involvement of the Ena/VASP family proteins – MENA, VASP, and EVL - in early synaptogenesis, we examined the effects of suppressing their activity on DF dynamics. We used peptides containing the FPPPP (FP4) repeat sequence from *Listeria monocytogenes* ActA protein (Niebuhr et al., 1997), which binds the EVH1 domains of MENA, VASP and EVL. Since the primary mode of activation of Ena/VASP proteins is through recruitment to their sites of action, sequestering them at mitochondria using FP4 fused to a mitochondrial targeting sequence (FP4-MITO) suppresses their activity (Bear et al., 2000). Additionally, we engineered an acute induction system by cloning these constructs into a doxycycline-inducible lentiviral expression vector. This allowed us to minimize harmful effects caused by long-term suppression of Ena/VASP (Dent et al., 2007; Kwiatkowski et al., 2007).

After 12 hours of doxycycline induction of mCherry-FP4-MITO expression, DF exhibited reduced overall dynamics and altered morphology, compared to the negative control mCherry-APPPP(AP4)-MITO (Fig. 1A). We examined the effect of FP4-MITO induction on DF tip motility, and found that average speed was substantially reduced compared to AP4-MITO (Fig. 1B). These data revealed that a greater proportion of DF from FP4-MITO-expressing neurons are, on average, “non-motile” (52.1%) during the duration of imaging, compared to AP4-MITO (29.8%) (Fig. 1C). Further, FP4-MITO significantly reduced the length of DF (Fig. 1D). These data suggest that the Ena/VASP family of AEFs influence DF morphology and motility.

**Figure 1:**
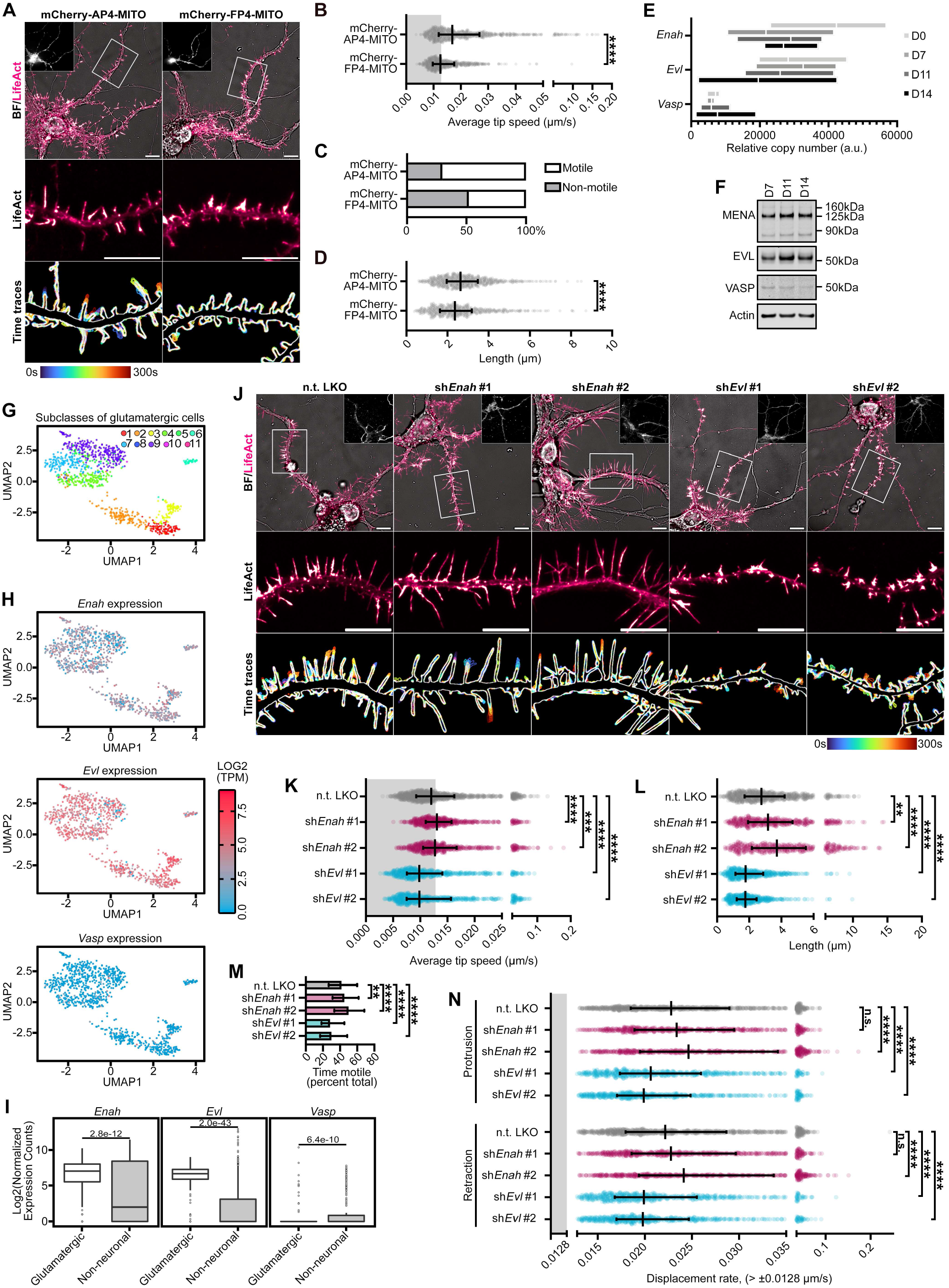
EVL is the predominant Ena/VASP paralog regulating dendritic filopodia. A. Live primary mouse cortical neurons at day *in vitro* 11 (D11) expressing EGFP-LifeAct and doxycycline-inducible mCherry-AP4-MITO (left column) or mCherry-FP4-MITO (right column), imaged 12 hours after doxycycline induction. Top: full cell image. Inset: mCherry-AP4/FP4-MITO expression. Middle: indicated segment of dendrite presented with an intensity-coded LUT. Bottom: maximum intensity projection of temporally color-coded binary mask outline, illustrating DF dynamics during imaging (5s interval, 5min duration). B. Scatterplot of average speed of DF tips, calculated as the average absolute tip displacement between successive timepoints. Grey shaded region indicates average speed less than 0.0128µm/s (non-motile DFs). Median +/- interquartile range (IQR). C. Bar graph of percent of total DF population with average tip speeds greater than 0.0128µm/s (motile) or less than 0.0128µm/s (non-motile). D. Scatterplot of average length reached during the duration of imaging. Median +/- IQR. E. RT-qPCR of RNA samples from primary mouse cortical neuron cultures at indicated days *in vitro*. C_T_s were normalized to the average of three housekeeping genes, and relative copy numbers were generated using 2^-ΔCT^ × 10^6^. Floating bars span the minimum and maximum data points, central line denotes mean. N = 3 biological replicates. F. Representative western blot of protein lysates from primary neuron cultures at indicated days *in vitro*, probed with antibodies targeting MENA, EVL, and VASP. G. Uniform manifold approximation and projection (UMAP) subcluster categories of single-cell RNA-seq expression profiles from mouse glutamatergic neurons. 1: L6b cortex (CTX), 2: L6 corticothalamic CTX, 3: L5 near-projecting CTX, 4: L6 intratelencephalic (IT) CTX, 5: L5 IT CTX, 6: L5 pyramidal tract CTX, 7: L2/3 IT CTX-1, 8: *Car3*, 9: L4/5 IT CTX, 10: L2/3 IT CTX *Ppp1r18*, 11: L2/3 IT CTX-2. H. Log_2_(TPM) expression in mouse of *Enah, Evl*, and *Vasp* across single-cell RNA-seq UMAP subcluster categories. I. Comparison of *Enah, Evl*, and *Vasp* expression in mouse across single-cell RNA-seq grouped by glutamatergic neurons versus non-neuronal cell types. J. Live D11 neurons expressing EGFP-LifeAct and pLKO-shRNA-TurboRFP targeting *Enah, Evl*, or non-targeting (n.t.) as indicated. Cells were transduced with shRNA lentiviral particles on D7. Top: full cell image. Inset: TurboRFP expression identifying shRNA-positive neurons. Middle: indicated segment of dendrite. Bottom: maximum intensity projection of temporally color-coded binary mask outline (5s interval, 5min duration). K. Scatterplot of average speed of DF tips. Grey shaded region indicates average speed < 0.0128µm/s (non-motile). Median +/- IQR. L. Scatterplot of average length reached during duration of imaging. Median +/- IQR. M. Bar graph of percent time motile (percent of time per DF in which instantaneous speed was greater than 0.0128µm/s). Median +/- IQR. N. Scatterplot of median protrusion and retraction rates of DFs (the median of values when instantaneous change in length was greater than +/-0.0128µm/s (motile)). Median +/- IQR. Statistical comparisons were made as follows: (B-D) Mann-Whitney test. n = 731-917, N = 3 biological replicates. (I) Welch’s t-test, n = 250 cells. (K-N) Kruskal-Wallis test and corrected for multiple comparisons. n = 630-1013, N = 3-5 biological replicates. * P < 0.05, ** P < 0.01, *** P < 0.001, **** P < 0.0001, n.s. is not significant. Scale bars = 10µm. See also Fig. S1.

We examined the expression of MENA, EVL and VASP in primary cortical neuron cultures throughout development *in vitro* by RT-qPCR. *Enah* and *Evl* expression was high during neuronal morphogenesis (D7) and during periods of early synaptogenesis (D11-D14) compared to *Vasp* (Fig. 1E). Western blot analysis corroborated the expression pattern at the protein level (Fig. 1F, Fig. S1E). Importantly, primary neuronal cultures inherently include a mixture of both glia and neurons; due to this unavoidable contamination by glial cells, it is difficult to accurately assess the individual contribution of each cell type to total mRNA or protein levels. To overcome this difficulty, we examined expression in excitatory cortical neurons specifically, by analyzing publicly-available mouse single-cell RNA-seq datasets from The Allen Brain Map data portal (Allen Cell Types Database (2015)). Glutamatergic cortical neurons from adult mouse (Fig. 1G-I) were confirmed to have strong expression of *Enah* and *Evl*, and comparatively low levels of *Vasp*. These expression patterns were also observed in adult human single-cell RNA-seq datasets (Fig. S1B-D; Allen Cell Types Database (2015)). Based on these analyses showing that MENA and EVL are the dominant Ena/VASP proteins in cortical neurons, we prioritized examining their role in DF.

To determine the influence of MENA and EVL on DF dynamics, we utilized shRNA targeting each paralog, introduced four days before experiments on D11, to limit potential impact on earlier neuronal morphogenesis (Fig. S1F,G). *Enah* knockdown (KD) DF retained a normal morphology, with significantly higher tip speed and length, compared to neurons expressing a non-targeting (n.t.) shRNA. In contrast, *Evl* KD neurons exhibited a stubby, flare-like DF morphology with profoundly suppressed dynamics (Fig. 1J-L, Video 1). In addition, *Enah* KD DF spent more of their time engaging in substantive motility (tip speed > 0.0128µm/s), while *Evl* KD DF exhibited significantly less time motile (Fig. 1M). To explore this effect on motility further, we examined the rate of protrusion and retraction as distinct events in control and KD cells. *Evl* KD DF had a slower rate of both protrusions and retractions compared with n.t. controls, while *Enah* KD DF overall trended towards faster rates (Fig. 1N). Together, these data implicate differential roles for MENA and EVL in DF dynamics, and identify EVL as a crucial regulator of DF morphology and motility.

### Tip enrichment of EVL precedes DF protrusion

To further investigate the function of MENA and EVL in regulating DF dynamics, we overexpressed EGFP-tagged constructs in primary cortical neuron cultures using lentivirus four days before assessing DFs on D11 (Fig. S2A). EGFP-MENA and EGFP-EVL were both enriched at the tips of protrusions (Fig. S2B), and exhibited a dose-dependent spectrum of phenotypes. Strong expression (signal-to-noise ratio (SNR)>1.5) of MENA had a high frequency of focal fan-like protrusions, while highly-expressing EVL neurons extended wide lamellipodia-like protrusions from the dendrites (Fig. S2B, arrowheads). These extreme phenotypes suggest unique functions for MENA and EVL in neuronal morphogenesis, and prompted us to restrict all subsequent quantification to low-expressing neurons (SNR<1.5). We profiled DF morphology using qualitative categories: normal, forked, multi-forked (>2 forks), flaring (lamellipodia-like protrusions), and complex forked + flaring morphology (Fig. S2C). We found that overexpressers of EVL and MENA exhibited the full range of morphologies observed in control neurons, with MENA increasing the proportion of multi-forked DF, and EVL promoting flaring. These findings further underscore the differential, non-overlapping function of MENA and EVL, and suggest that each paralog promotes distinct filopodia-like structures in neurons.

Expression of MENA and EVL each enhanced overall DF dynamics, tip speed, and protrusion rate, though DF length was not significantly altered under any conditions (Fig. 2A-E). In both low and high-expressing neurons, EGFP-MENA and EGFP-EVL were readily visible at DF tips, compared to the cytosolic localization of EGFP alone (Fig. 2A, S2B, Video 2). This tip enrichment was dynamic throughout the lifetime of a DF, with EVL exhibiting a higher variance than MENA in mean intensity levels at a tip ROI (Fig. 3A,B), indicating that EVL tip localization is more dynamic compared to MENA. Examination of the relationship between tip enrichment and motility by kymography revealed that while EGFP alone remained cytosolic and uniform during DF protrusion and retraction, EGFP-EVL consistently exhibited distinct and persistent enrichment at the DF tip starting before a protrusion event. In contrast, EGFP-MENA was enriched at the tip during protrusion in only a subset of DF (Fig. 3C).

**Figure 2:**
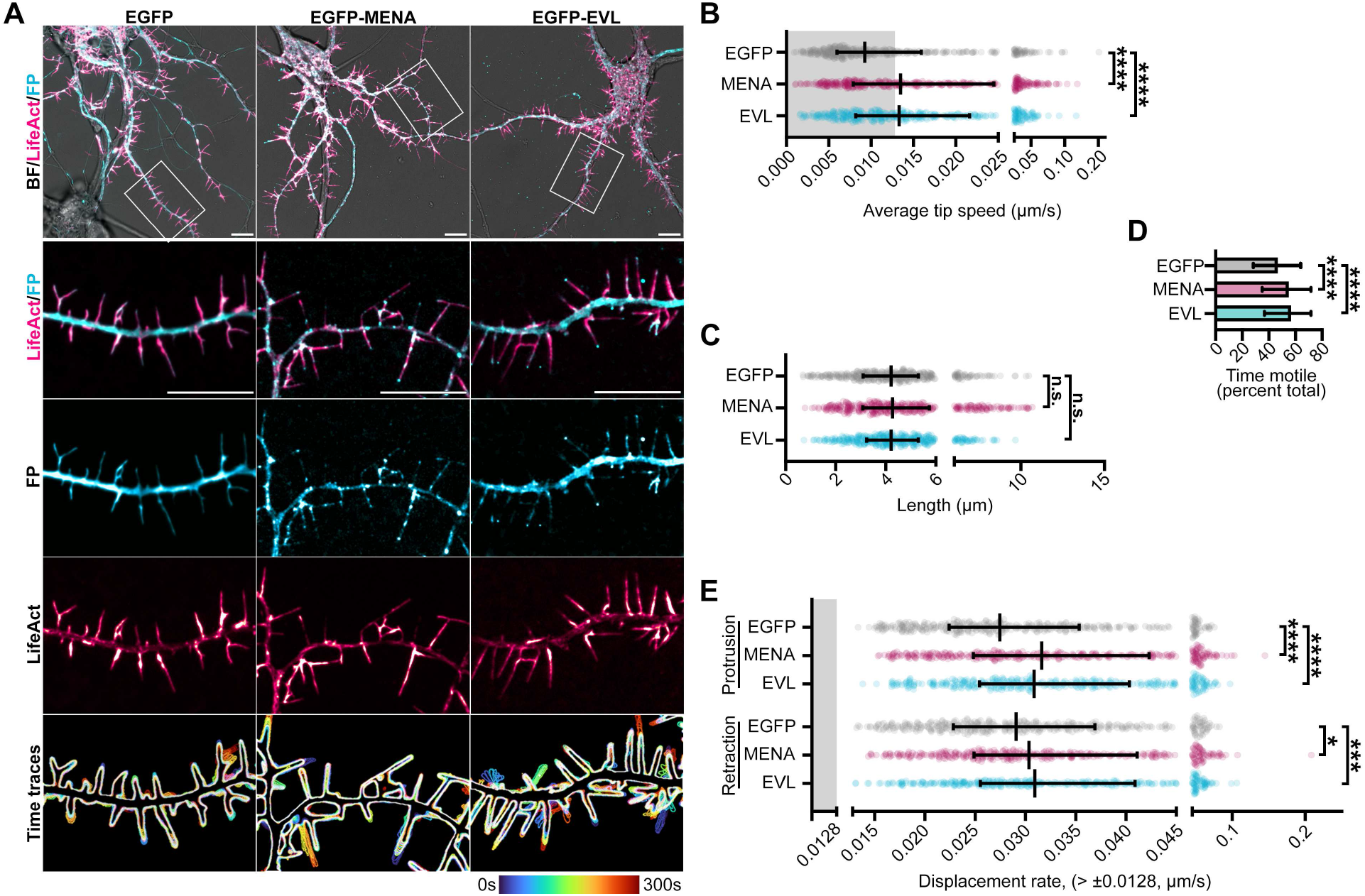
MENA and EVL overexpression enhance DF motility. A. Live primary mouse cortical neurons at day *in vitro* 11 (D11) expressing mRuby2-LifeAct and EGFP (left column), EGFP-MENA (middle column), or EGFP-EVL (right column). Row 1: full cell image. Row 2: indicated segment of dendrite. Row 3 and 4: individual fluorescent channels presented with intensity-coded LUTs. Row 5: maximum intensity projection of temporally color-coded binary mask outline, illustrating DF dynamics during imaging (5s interval, 5min duration). B. Scatterplot of average speed of DF tips, calculated as the average absolute tip displacement between successive timepoints. Grey shaded region indicates average speed less than 0.0128µm/s (non-motile DFs). Median +/- Interquartile range (IQR). C. Scatterplot of average length reached during the duration of imaging. Median +/- IQR. D. Bar graph of percent time motile (percent of time per DF in which instantaneous speed was greater than 0.0128µm/s). Median +/- IQR. E. Scatterplot of median protrusion and retraction rates of DFs (the median of values when instantaneous change in length was greater than +/-0.0128µm/s (motile)). Median +/- IQR. Statistical comparisons of EGFP vs. MENA vs. EVL (B-E) were made by Kruskal-Wallis test and corrected for multiple comparisons. n = 368-389, N = 3 biological replicates. Scale bars = 10µm. * P < 0.05, ** P < 0.01, *** P < 0.001, **** P < 0.0001, n.s. is not significant. See also Fig. S2, Supp. Video 2.

**Figure 3:**
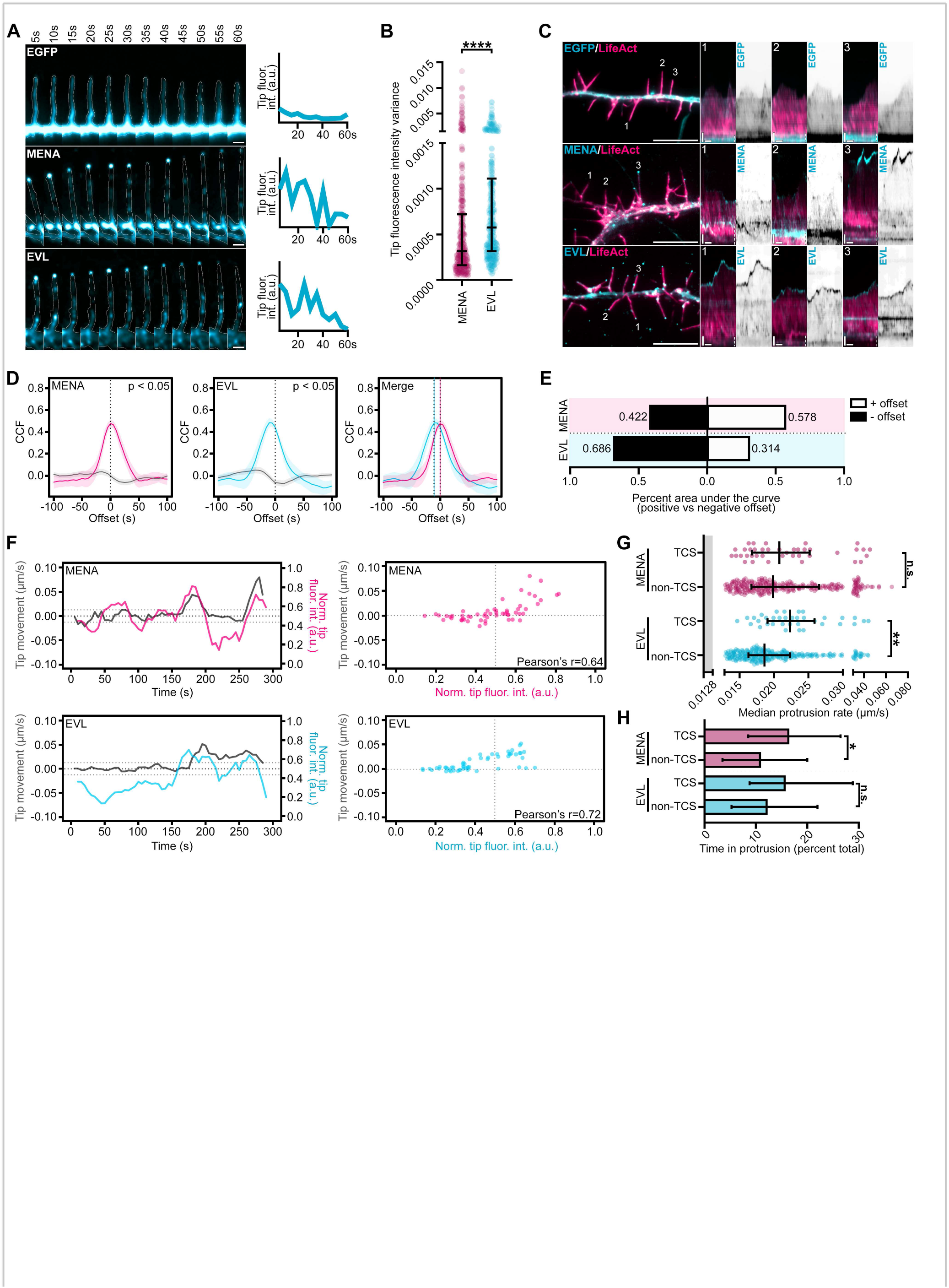
Tip enrichment of EVL precedes DF protrusion. A. Filmstrip of live primary mouse cortical neurons at day *in vitro* 11 (D11) expressing EGFP (top row), EGFP-MENA (middle row), or EGFP-EVL (bottom row) in a representative DF showing localization dynamics presented with an intensity-coded LUT. Line scan of normalized fluorescence intensity over time at tracked tip (right; 5s interval, 1min duration). Scale bar = 1µm. B. Scatterplot of tip fluorescence variance at DF tips normalized to local background demonstrates range of tip enrichment per DF and hence magnitude of on-off dynamics. Median +/- interquartile range (IQR). C. Live D11 neurons expressing mRuby2-LifeAct and EGFP (top row), EGFP-MENA (middle row), or EGFP-EVL (bottom row). Segment of dendrite (left).; scale bar = 10µm. Numbers indicate DF position analyzed by kymograph (right), highlighting protein localization dynamics during DF motility (5s interval, 5min duration). Vertical scale bar = 1µm, horizontal scale bar = 1min, dashed line indicates dendrite. D. Line plots of cross-correlation function (CCF) of normalized tip fluorescence intensity and tip motility as a function of time offset for top-correlating subcluster (TCS; color lines) versus non-TCS (grey lines) determined in Fig. S3B, of DFs from D11 neurons expressing EGFP, EGFP-ENAH, or EGFP-EVL. Peak CCF at negative offset values indicates that fluorescence enrichment precedes motility, while peak CCF at positive offset values indicates fluorescence enrichment follows motility. Mean +/- 95% CI. Peak CCF GFP = 0.43 at offset 0, .413 at offset -5s; ENA = 0.475 at offset 0, 0.464 at offset +5s; EVL =, 0.482 at offset -10s, 0.480 at offset -5s. Statistical significance determined by peak CCF > 2/√(n-|offset|). E. Percent of area under the curve at positive and negative offset values for indicated conditions. F. Line plots (left) and scatterplots (right) for one representative TCS DF from D11 neurons from indicated conditions. Left: line plot of tip motility (grey, left y-axis) and normalized tip fluorescence (color, right y-axis) during imaging (5s interval, 5min duration). Right: scatterplot of normalized tip fluorescence and tip motility at individual timepoints demonstrating strength of relationship by Pearson’s correlation test. G. Scatterplot of median protrusion rates for TCS versus non-TCS DFs from indicated conditions. Protrusion rate is the median of values when instantaneous change in length was greater than +0.0128µm/s (motile, protruding). Median +/- interquartile range (IQR). H. Bar graph of percent of time in protrusion during the duration of imaging for TCS versus non-TCS DFs from indicated conditions. Percent time in protrusion is calculated as the percent of time per DF in which positive change in length between successive timepoints was greater than 0.0128µm/s. Median +/- IQR. Statistical comparisons were made as follows: (B) Mann-Whitney test. n = 368-389, N = 3 biological replicates. (G,H) Kruskal-Wallis test and corrected for multiple comparisons. n = 360-386, TCS n = 42 (MENA), 39 (EVL), non-TCS n = 320 (MENA), 321 (EVL). N = 3 biological replicates. * P < 0.05, ** P < 0.01, *** P < 0.001, **** P < 0.0001, n.s. is not significant. See also Fig. S3.

To determine whether tip enrichment of MENA or EVL is correlated with DF protrusive behavior, we utilized the filopodia analysis program, Filopodyan (Urbančič et al., 2017). Filopodyan cross-correlates tip fluorescence and motility in time, determines the time offset at which correlation is highest, and identifies, using hierarchical clustering, the top-correlating subcluster (TCS) of DFs in each condition. This enables examination of DF motility within the TCS compared to DFs with poor correlation of tip fluorescence and motility (non-TCS). Not surprisingly, without clustering and by including the entire population of tracked DF in our analysis, EGFP-MENA or EGFP-EVL tip fluorescence and motility were not significantly correlated, in contrast to the TCS populations for each condition (Fig. S3A-C). Cross-correlation function (CCF) revealed that the TCS of EGFP-EVL DFs exhibited highest correlation at offsets of -10s and -5s, indicating that EVL tip enrichment precedes motility. In contrast, CCF in EGFP-MENA DFs peaked at 0s offset, but skewed towards positive offset, suggesting that more TCS DFs exhibited tip enrichment with or following motility (Fig. 3D-F).

For further characterization of the TCS populations, we compared the protrusive motility in TCS versus non-TCS DFs from each condition. Tip enrichment was associated with increased rates of protrusion in EGFP-EVL TCS DF, while EGFP-MENA TCS DF spent more time engaging in protrusive events (Fig. 3G,H). To validate the TCS clustering method, 1000 randomized datasets were generated, and identical cross-correlation analysis was performed on datasets with similar subcluster size (Fig. S3D). At -10s offset, TCSs from randomized datasets of EGFP-EVL DFs showed significantly lower cross-correlation than the observed TCS (bootstrap p = 0.003), while the TCS of EGFP-MENA DFs at 0s offset were not significantly different than randomized data (bootstrap p = 0.114; Fig. S3E). Together, these data suggest that although MENA overexpression may promote DF dynamics, EVL tip enrichment uniquely predicts protrusion events, providing further evidence for a distinct function for EVL in DF motility.

### The EVH1 domain is required for EVL localization and activity

To investigate the direct involvement of EVL in DF dynamics, we examined primary cortical neurons derived from EVL knockout mice (EVL KO; (Kwiatkowski et al., 2007)), and compared them to wild-type (WT) neurons throughout a developmental time course *in vitro*. EVL KO neurons underwent normal overall morphogenesis, and developed primary neurites, axons, and branching comparable to WT neurons (Fig. S4A-E). Importantly, no compensatory upregulation of MENA or VASP was observed in EVL KO cultures (Fig. S4F,G). EVL KO neurons manifested increasingly pronounced defects in DF morphology and dynamics over time, and by D9 the majority of dendrites exhibited very short filopodia or small lamellipodia-like flares (Fig. 4A, Video 3), with maximal reductions in speed, length, and protrusion rate (Fig. 4B-C, Fig. S4H). This suggests that EVL is the dominant protein regulating DF dynamics after D9, or that sustained loss of EVL during DF morphogenesis drives these phenotypes.

**Figure 4:**
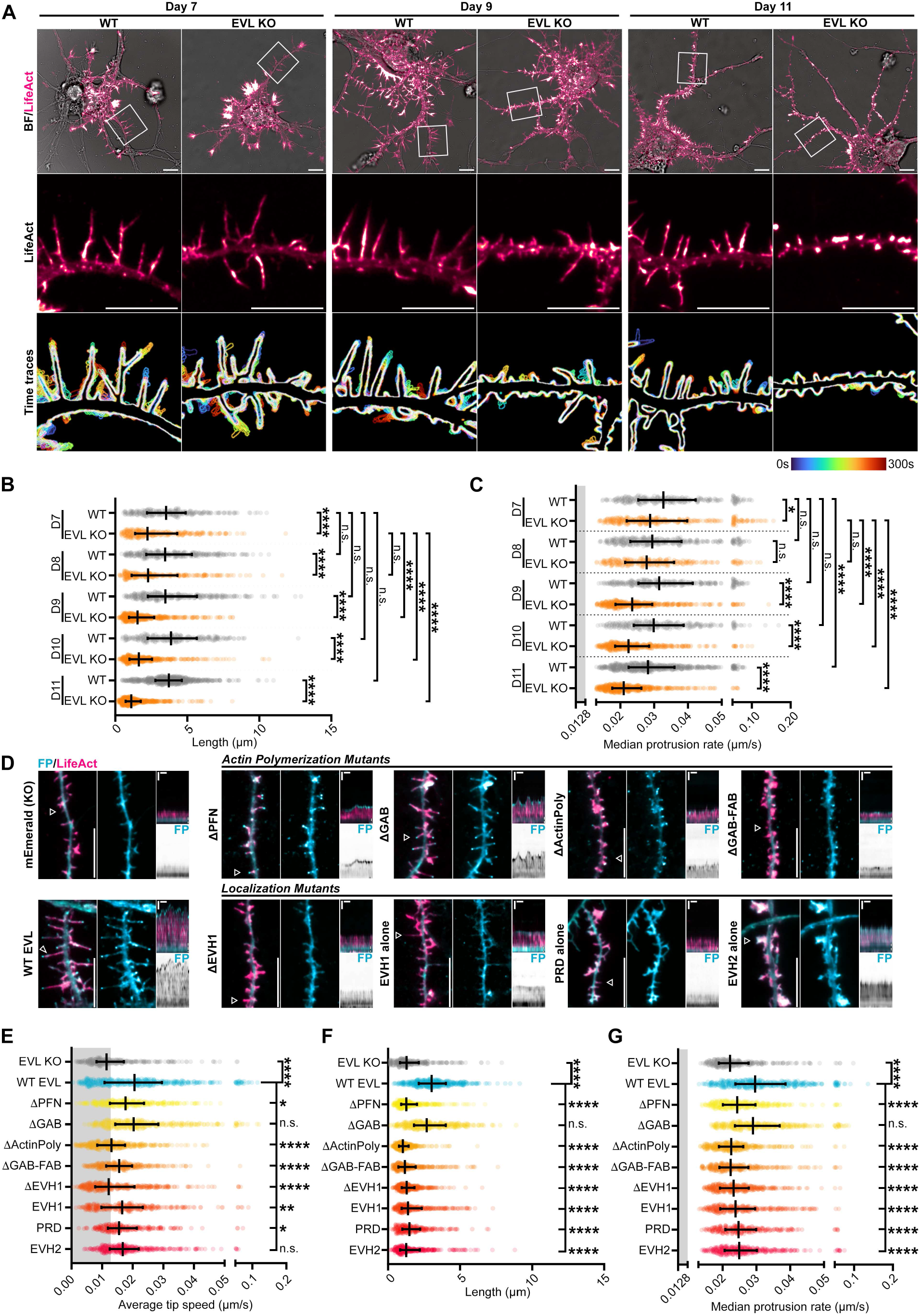
EVL regulates DF morphogenesis and motility through membrane-targeted actin polymerization. A. Live primary cortical neurons derived from wildtype (WT) or EVL knockout (KO) mice at indicated days *in vitro* expressing mRuby2-LifeAct. Top: full cell image. Middle: indicated segment of dendrite presented with an intensity-coded LUT. Bottom: maximum intensity projection of temporally color-coded binary mask outline, illustrating DF dynamics during imaging (5s interval, 5min duration). Scale bars = 10µm. B. Scatterplot of average length reached during the duration of imaging for indicated conditions. Median +/- interquartile range (IQR). C. Scatterplot of median protrusion rates of DFs (the median of values when instantaneous change in length was greater than +0.0128µm/s (motile), protruding). Median +/- IQR. D. Segment of dendrite from live EVL KO cortical neurons at D11 expressing mRuby2-LifeAct and mEmerald alone, or mEmerald-tagged wildtype EVL or mutants of EVL as indicated (left). Middle: mEmerald fluorescence intensity presented with an intensity-coded LUT. Right: kymograph of DF position indicated by arrowhead in column 1 (5s interval, 5min duration). Vertical scale bar = 1µm, horizontal scale bar = 1min, dashed line indicates dendrite. E. Scatterplot of average speed of DF tips, calculated as the average absolute tip displacement between successive timepoints. Grey shaded region indicates average speed less than 0.0128µm/s (non-motile DFs). Median +/- IQR. F. Scatterplot of average length reached during the duration of imaging for indicated conditions. Median +/- IQR. G. Scatterplot of median protrusion rates of DFs for indicated conditions. Median +/- IQR. Statistical comparisons were made as follows: (B-C) Kruskal-Wallis test and corrected for multiple comparisons. n = 288-641, N = 3 biological replicates. (E-G) made by Kruskal-Wallis test and corrected for multiple comparisons. n = 248-397, N = 3-5 biological replicates. * P < 0.05, ** P < 0.01, *** P < 0.001, **** P < 0.0001, n.s. is not significant. See also Fig. S4, Supp. Video 3,4.

In order to determine the mechanism of EVL’s regulation of DF dynamics, we reconstituted either WT mEmerald-EVL, or mutants of EVL, in KO neurons by lentiviral transduction at D7 *in vitro* (Fig. S4I,J). Re-introduction of WT EVL fully rescued DF motility, length, and tip localization (Fig. 4D-G, Fig. S4K, Video 4). We investigated two categories of EVL mutants: domain deletions that have been described to impact actin polymerization, and deletions that impact EVL localization. EVL polymerizes actin through direct interactions with profilin (PFN), G-actin, and F-actin (Chereau and Dominguez, 2006). Reconstitution of a mutant lacking the PFN-binding site (ΔPFN) showed tip localization and slow DF elongation, while a G-actin binding mutant (ΔGAB) resulted in a full rescue of tip enrichment, motility, and length. In contrast, loss of both profilin- and G-actin-binding (ΔActinPoly), or both the G- and F-actin-binding domains (ΔGABFAB) resulted in DF with dynamic tip localization, but low motility (Fig. 4D-G, Fig. S4K, Video 4).

Targeted localization of Ena/VASP proteins is critical to their function. The EVH1 domain interacts with proline-rich residues on binding partners, and the proline-rich domain (PRD) contains several proline-rich motifs that bind SH3 domains in addition to profilin (Lambrechts et al., 2000; Reinhard et al., 1995). Deletion of the EVH1 domain (ΔEVH1) failed to rescue DF, while expression of the EVH1 domain alone showed a diffuse tip enrichment, with partial restoration of dynamics but not length. Expression of the PRD alone caused slightly enhanced motility and altered morphology. Lastly, expression of the EVH2 domain alone, which contains the GAB and FAB domains, as well as a coiled-coiled region important for Ena/VASP tetramerization, promoted the formation of large lamellipodia-like flares and diffuse localization throughout the protrusion (Fig. 4D-G, Fig. S4K, Video 4). Together, these results indicate that the EVH1 domain is required for EVL’s enrichment to DF tips, while actin polymerization, primarily mediated by the profilin-binding region, is required for promoting length and motility of DF.

### EVL tip localization is necessary and sufficient for DF motility

Since localized recruitment is the central mechanism by which Ena/VASP proteins are activated, we employed an optogenetic approach, iLID (improved light-induced dimer, (Guntas et al., 2015; Zimmerman et al., 2016)), to examine the effects of acutely altering the localization of EVL on DF dynamics. iLID is a LOV2-based technology, in which each half of a heterodimerizing protein pair (SsrA-SspB) is fused to (1) a protein of interest, or (2) the dark state-obscured C-terminus of the photosensitive protein *As*LOV2 along with a plasma membrane targeting sequence (CAAX) or mitochondrial targeting sequence (MITO). During exposure to 488nm light, LOV2 undergoes a conformational change, removing steric occlusion of SsrA to permit its dimerization with SspB, and thereby inducibly and reversibly recruiting the protein of interest to the plasma membrane or to the mitochondria (Fig. 5A). We generated EVL-iLID and expressed it together with CAAX-iLID or MITO-iLID in order to acutely localize EVL to or away from sites of action, respectively, during photostimulation (Fig. 5B). In glial cells (used for design validation), plasma membrane recruitment of EVL-iLID resulted in increased leading edge and filopodial tip enrichment of EVL, and enhanced lamellipodial and filopodial protrusion (Fig. 5B, S5A). In contrast, photorecruitment of EVL-iLID to mitochondria resulted in reduced membrane localization, and suppression of membrane dynamics (Fig. 5B, S5B). An important caveat is that iLID does not alter the activity of EVL; therefore, high expression of EVL-iLID exhibits similar effects to EVL overexpression, even in the absence of photostimulation. To minimize this confounding factor, we examined low expressers of EVL-iLID (SNR<1.5) reconstituted in EVL KO neurons.

**Figure 5:**
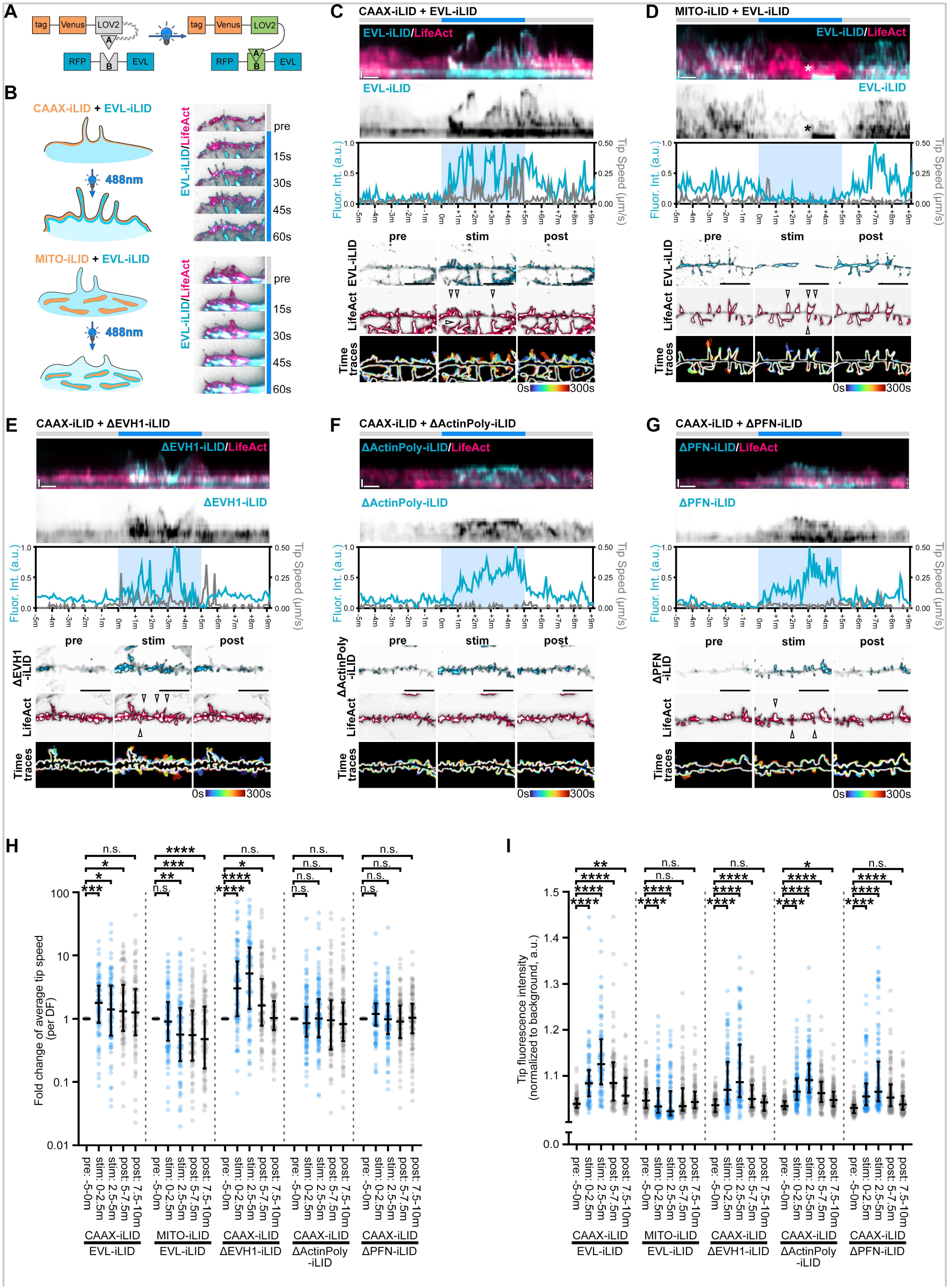
EVL tip localization is necessary and sufficient for DF motility. A-B Schematic of optogenetic EVL-iLID system. (A) The SsrA peptide (A) is embedded in the Jɑ helix of *As*LOV2, and its binding partner protein, SspB (B), is tagged to EVL. LOV2-SsrA is additionally tagged by a plasma membrane targeting motif (CAAX) or mitochondrial targeting sequence (MITO). Upon irradiation with 488nm light, the Jɑ helix unwinds and reveals the occluded SsrA peptide, enabling dimerization with SspB. (B) Each iLID component – EVL-iLID and either CAAX-iLID or MITO-iLID, are introduced to target cells by lentivirus. Exposure of the whole cell to 488nm light recruits EVL to the plasma membrane or to the mitochondria, and influences actin-based protrusions including lamellipodia and filopodia (filmstrip, right). C-G. Segment of dendrite from live primary cortical neurons derived from EVL KO mice at D11 expressing iRFP670-LifeAct, and indicated iLID constructs. (C) WT EVL-iLID with CAAX-iLID. (D) WT EVL-iLID with MITO-iLID. (E) ΔEVH1-iLID with CAAX-iLID. (F) ΔActinPoly-iLID with CAAX-iLID. (G) ΔPFN-iLID with CAAX-iLID. Neurons were photostimulated with 488nm light for 5min, and imaged for 5min before, during, and after photostimulation. Rows 1-2: Kymograph of representative DF position. 15min duration, 5s interval, vertical scale bar = 1µm, horizontal scale bar = 1min, dashed line indicates dendrite. Row 3: Line plot of normalized fluorescence intensity (cyan) and DF tip speed (grey) over time. Rows 4-5: segment of dendrite showing localization of EVL-iLID and LifeAct at single timepoint during photostimulation experiment. Individual channels displayed with black subtraction for ease of morphological comparison. Scale bars = 10µm. Row 6: maximum intensity projection of temporally color-coded binary mask outline. Arrowheads indicate DFs which exhibited altered dynamics following photostimulation. Asterisk in (D) indicates a mitochondrion that moved into the kymograph. H. Fold change in average tip speed of 2.5min bins during or post-photostimulation for indicated iLID conditions, relative to average tip speed pre-photostimulation. Median +/- interquartile range (IQR). I. Scatterplot of background-normalized fluorescence intensity of EVL-iLID pre-photostimulation, or in 2.5min bins during or post-photostimulation, for indicated iLID conditions. Median +/- IQR. All statistical comparisons were paired and followed individual DFs during phases of stimulation, using Friedman’s Test. n = 85-120, N = 3 biological replicates. * P < 0.05, ** P < 0.01, *** P < 0.001, **** P < 0.0001, n.s. is not significant. See also Fig. S5, Supp. Video 5.

In CAAX-iLID neurons, photostimulation resulted in the rapid recruitment of EVL-iLID to the plasma membrane and to DF tips, concurrent with DF elongation and increased motility; these activities subsided after termination of light exposure (Fig. 5C,H,I, Video 5). When co-expressed with MITO-iLID, EVL-iLID was pulled away from DF tips and to mitochondria in the dendrite during photostimulation, resulting in reduced motility, which recovered only partially during the observation period following light withdrawal (Fig. 5D,H,I, Video 5). Together, these data demonstrate that EVL localization at the tips of DF is both necessary and sufficient for DF motility. Further, we used our CAAX-iLID/EVL-iLID system to acutely manipulate the localization of the ΔEVH1 mutant to the membrane. Expression of mEmerald-tagged ΔEVH1 in EVL KO neurons did not enhance DF dynamics or promote tip enrichment (Fig. 4). However, upon photostimulation, ΔEVH1-iLID became tip-enriched, and substantially increased DF motility, which subsided after termination of light exposure (Fig. 5E,H,I, Video 5). Conversely, the ΔActinPoly-iLID mutant, which lacks the ability to polymerize actin, exhibited membrane enrichment but no increase in DF motility during light exposure, while ΔPFN-iLID, which polymerizes actin very weakly, showed enrichment with a modest but non-significant increase in motility (Fig. 5G-I, Video 5). These data show that although the EVH1 domain is required for EVL localization and DF motility, engineered membrane recruitment using CAAX-iLID rescues KO phenotypes during transient photostimulation.

### The I-BAR protein MIM/MTSS1 is a binding partner of EVL

To further investigate the mechanism by which EVL regulates DF, we took an unbiased quantitative proteomics approach to identify putative protein partners of EVL that co-regulate DF motility. For that purpose, we performed affinity purification mass spectrometry (AP-MS) experiments using endogenous EVL as bait in lysates from cultured cortical neurons (Fig. S6A). Immunoprecipitated proteins were separated by SDS-PAGE and the interactomes were subjected to label-free quantification. Spectrum counting of EVL immunoprecipitates versus non-immune serum negative control immunoprecipitates uncovered a high-confidence putative interaction with MIM/MTSS1, and confirmed interactions with *bona fide* Ena/VASP binding partners including profilin isoforms (Fig. S6B). MIM is an Inverse BAR (I-BAR) domain protein, which binds and deforms the plasma membrane at sites of PIP2 enrichment to promote outward membrane curvature (Mattila et al., 2007; Saarikangas et al., 2009). In addition, MIM has been shown to promote Arp2/3-dependent assembly of branched actin networks (Lin et al., 2005). Together, these activities promote the generation of a proto-protrusion, a critical first step in DF initiation in cortical neurons (Saarikangas et al., 2015). Therefore, we prioritized studying MIM as an EVL binding partner in our expansion of the mechanism by which EVL regulates DF motility.

We confirmed the interaction between EVL and MIM by performing the reciprocal co-IP with tagged plasmid proteins in HEK293T, demonstrating that EVL robustly co-IPs with MIM (Fig. 6A). To determine the domains involved in the EVL-MIM interaction, we expressed MIM with full-length EVL or the ΔEVH1 mutant. Although MIM interacted with full-length EVL, the interaction was lost between ΔEVH1 and MIM, suggesting that the EVH1 domain is important for MIM-EVL interaction (Fig. 6B).

**Figure 6:**
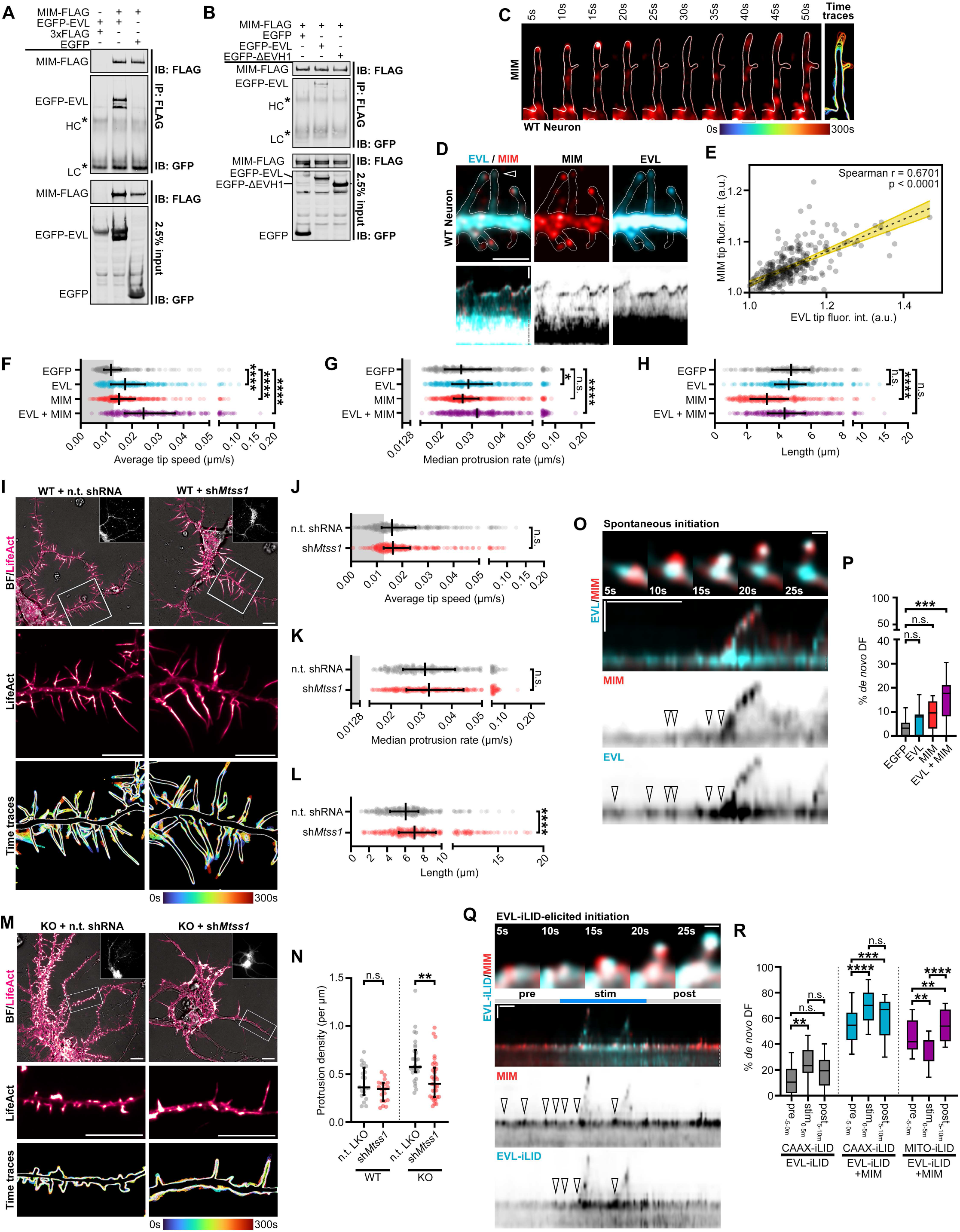
MIM/MTSS1 cooperates with EVL to promote DF initiation and motility. A-B. Representative western blots of co-immunoprecipitation experiments confirming reciprocal interaction of MIM with EVL (A) and dependence of interaction on EVL’s EVH1 domain (B). HEK293T overexpressing indicated constructs were lysed and immunoprecipitated (IP) with FLAG antibody to pull down MIM-FLAG, resolved by SDS-PAGE, and immunoblotted (IB) with indicated antibodies. C. Filmstrip showing localization dynamics of MIM-mRuby2 during protrusive motility in a WT cortical neuron DF at day *in vitro* 11 (D11; left), and maximum intensity projection of temporally color-coded binary mask outline (right; 5s interval, 1m duration). Scale bar = 1µm. D. Segment of dendrite from a WT cortical neuron at D11 co-expressing mEmerald-EVL and MIM-mRuby2 (top;), and kymograph of DF indicated by arrowhead (bottom; 5s interval, 5min duration. Dendrite segment: Scale bar = 5µm. Kymographs: Horizontal scale bar = 1min, vertical scale bar = 1µm, dashed line indicates dendrite. E. Scatterplot of background-normalized EVL and MIM tip fluorescence at individual timepoints. Strong correlation observed by Spearman’s correlation test. F. Scatterplot of average speed of WT DF tips for indicated conditions, calculated as the average absolute tip displacement between successive timepoints. Grey shaded region indicates average speed less than 0.0128µm/s (non-motile DFs). Median +/- interquartile range (IQR). G. Scatterplot of median protrusion rates of WT DFs for indicated conditions (the median of values when instantaneous change in length was greater than +0.0128µm/s (motile, protruding)). Median +/- IQR. H. Scatterplot of average length of WT DFs reached during the duration of imaging for indicated conditions. Median +/- IQR. I. WT neurons expressing EGFP-LifeAct and pLKO-shRNA-TurboRFP targeting *Mtss1* or non-targeting as indicated. Cells were transduced with shRNA lentiviral particles on D7. Top: full cell image. Inset: TurboRFP expression identifies shRNA-positive neurons. Middle: indicated segment of dendrite. Bottom: maximum intensity projection of temporally color-coded binary mask outline (5s interval, 5min duration). Scale bars = 10µm. J. Scatterplot of average speed of DF tips for indicated conditions. Median +/- IQR. K. Scatterplot of median protrusion rates of DFs for indicated conditions. Median +/- IQR. L. Scatterplot of average length of DFs reached during the duration of imaging for indicated conditions. Median +/- IQR. M. EVL KO neurons expressing EGFP-LifeAct and pLKO-shRNA-TurboRFP targeting *Mtss1* or non-targeting as indicated. Top: full cell image. Inset: TurboRFP expression identifies shRNA-positive neurons. Middle: indicated segment of dendrite. Bottom: maximum intensity projection of temporally color-coded binary mask outline (5s interval, 5min duration). Scale bars = 10µm. N. Scatterplot of protrusion density along dendrites from indicated conditions. Protrusions were defined as any actin-rich proturbences visibly extending beyond the dendrite. Median +/- IQR. O. Example of spontaneous DF initiation. Row 1: filmstrip indicating localization of mEmerald-EVL and MIM-mRuby2 during DF initiation in a D11 WT neuron. Scale bar = 0.5µm. Rows 2-4: kymograph of mEmerald-EVL and MIM-mRuby2 during initiation. Arrowheads indicate foci of protein enrichment preceding initiation or motility events. 5s interval, 5min duration, horizontal scale bar = 1min, vertical scale bar = 1µm, dashed line indicates dendrite. P. Box-and-whisker plot of newly initiated DFs as a percent of total DFs during the duration of imaging. Median +/- IQR. Q. Example of EVL-iLID-elicited DF initiation. Row 1: filmstrip indicating localization of EVL-iLID and MIM-iRFP670 during photostimulation in a D11 KO neuron. Scale bar = 0.5µm. Rows 2-4: kymograph of EVL-iLID and MIM-iRFP670 during initiation. Arrowheads indicate foci of protein enrichment preceding initiation or motility events. 5s interval, 15min duration, horizontal scale bar = 1min, vertical scale bar = 1µm, dashed line indicates dendrite. R. Box-and-whisker plot of newly initiated DFs as a percent of total DFs during the duration of imaging before, during, and following iLID stimulation with indicated constructs. Statistical comparisons were made as follows: (F-H) Kruskal-Wallis test and corrected for multiple comparisons. n = 273-345, N = 3 biological replicates. (J-L) Mann-Whitney test. n = 362-363, N = 3 biological replicates. (N) made by Mann-Whitney test. n = 19-35 dendrites, N = 3 biological replicates. (P,R) Kruskal-Wallis test and corrected for multiple comparisons. n = 9-16, N = 3 biological replicates.* P < 0.05, ** P < 0.01, *** P < 0.001, **** P < 0.0001, n.s. is not significant. See also Fig. S6, Supp. Videos 6,7.

### MIM/MTSS1 cooperates with EVL to promote DF initiation and motility

In neurons, expression of MIM was robust throughout *in vitro* development and was not affected by EVL KO (Fig. S6C). To investigate the relationship between MIM and EVL in regulating DF, we expressed MIM alone, EVL alone, or both MIM and EVL in WT cortical neurons and examined DF motility. When MIM was expressed alone in developing WT neurons, it was enriched at DF tips during protrusive motility (Fig. 6C) and increased absolute DF tip dynamics; nonetheless, MIM expression was not sufficient to significantly increase protrusion rate (Fig. 6F,G). When MIM is co-expressed with EVL, the two proteins strongly co-localized at the DF tip (Fig. 6D,E) and together they dramatically enhanced all aspects of motility compared to control and to EVL-expressing neurons (Fig. 6F,G, Video 6). Interestingly, MIM expression significantly decreased DF length, a phenotype that was ameliorated by EVL co-expression (Fig. 6H). These findings suggest that the relationship between MIM and EVL is cooperative, and co-expression manifests additive phenotypes compared to expression of MIM or EVL alone.

Furthermore, when expressed alone in KO neurons, MIM robustly localizes to the tips of short DFs and protrusions; these sites of enrichment fail to elongate or exhibit substantial protrusive motility (Fig. S6E,G-I). Intriguingly, whereas MIM and WT EVL co-expression promotes a strong rescue of DF morphology and motility in KO neurons, co-expression of MIM and ΔEVH1 resulted in only a moderate —yet significant— intermediate phenotype, with diffuse ΔEVH1 localized proximally to the MIM-enriched DF tip (Fig. S6F). These observations underscore the necessity of the EVH1 domain for EVL’s tip localization, while suggesting that ΔEVH1 EVL, which is able to polymerize actin, can partially restore EVL activity when MIM is abundant.

To determine if MIM is competent to promote dynamics at the tip, we generated MIM-iLID to acutely manipulate MIM localization. Light-stimulated recruitment of MIM-iLID to the membrane failed to promote initiation or DF motility in WT neurons (data not shown). This may be due to MIM’s requirement of upstream regulation by PIP2 for its activity, or saturation of suitable sites (Mattila et al., 2007). In contrast, optogenetic sequestration of MIM-iLID at mitochondria reduced lamellipodia dynamics in glial cells, and resulted in modest but significant reductions in DF motility in WT neurons (Fig. S6J-L). This suggests that tip-localized MIM facilitates actin polymerization.

To examine the requirement of MIM in regulating DF, we knocked-down MIM using *Mtss1*-directed shRNA (Fig. 6J-K and S6D). MIM KD in WT neurons increased DF length (Fig. 6L), which is the opposite phenotype observed in MIM overexpressers (Fig. 6H) and in agreement with previous findings (Galbraith et al., 2018). However, in contrast to acute inhibition of MIM using the iLID system, MIM KD did not alter DF dynamics (Fig. 6J-K). These results suggest that MIM is not required for the motility of DF, and in fact, it limits DF length. We examined the effects of MIM KD on DF in the absence of EVL. MIM KD in EVL KO neurons resulted in a substantial reduction in the density of actin-rich protrusions along the dendrite, while in WT neurons, no differences in protrusion density were observed between *Mtss1* KD and control (Fig. 6I,M,N). These findings suggest that in the absence of both EVL and MIM, DF initiation is compromised.

We further investigated the relationship between MIM and EVL in early DF initiation events. During spontaneous DF initiation, both EVL and MIM exhibit tip localization during protrusion, and are strongly coincident preceding an initiation event. Visualization of the formation of a proto-protrusion on the dendrite by kymography suggests that MIM enrichment is a requisite event for DF initiation, but membrane enrichment alone is not sufficient for productive initiation of a DF (Fig. 6O, arrowheads). Further, co-expression of MIM and EVL promoted a significant increase in *de novo* DF initiation compared to expression of either protein alone (Fig. 6P). To further examine the cooperation between EVL and MIM, we examined initiation events elicited by photorecruitment of EVL-iLID to the membrane by CAAX-iLID. Neurons co-expressing EVL-iLID and MIM-iRFP exhibited extremely dynamic and unstable DF, with more initiation events at baseline before photostimulation, compared to EVL-iLID alone (Fig. 6Q,R, Video 7). Following photostimulation, initiation was further increased with CAAX-iLID membrane recruitment, and subsided after stimulation is suspended (Fig. 6R). Importantly, forward protrusion was not observed unless both EVL and MIM are present in the proto-protrusion, which is consistent with the spontaneous DF initiation results (Fig. 6Q, arrowheads). In addition, we used MITO-iLID to sequester EVL-iLID away from the membrane; photostimulation in these experiments suppressed DF initiation, which robustly recovered after stimulation was suspended (Fig. 6R, Video 7). These data suggest that, while EVL promotes actin elongation that is crucial for generating protrusive force, MIM is required to “license” the initiation of a nascent DF by signaling for actin nucleation.

### Arp2/3 and formins differentially contribute to DF initiation and motility

I-BAR proteins, such as MIM, promote the activation of Arp2/3-mediated actin nucleation (Lin et al., 2005; Saarikangas et al., 2015), which plays a significant role in building a branched actin architecture in DF (Korobova and Svitkina, 2010). The prevailing model of actin remodeling in DF proposes that Arp2/3 nucleates branched actin filaments throughout the DF length, which are elongated by formins at the tip to control motility (Hotulainen et al., 2009). To explore how Arp2/3 and formin activity interface with EVL during DF dynamics, we investigated the effects of their acute pharmacological inhibition using CK-666 (Nolen et al., 2009) and the pan-formin inhibitor SMIFH2 (Rizvi et al., 2009), respectively, in WT and EVL KO neurons. Neurons were imaged for 5 minutes before and 40 minutes after treatment, and assessed for changes in motility and actin dynamics throughout the duration of imaging. We examined growth cone dynamics as a read-out for approximating the time to maximal inhibition of Arp2/3 (∼20 minutes) and formin activity (∼25 minutes) in neurons.

Following Arp2/3 inhibition, WT DF exhibited a striking increase in motility and rapid elongation compared to DMSO controls (Fig. 7A,C, Video 8), in agreement with previous studies (Galbraith et al., 2018; Hotulainen et al., 2009; Spence et al., 2016). This behavior is likely due to a reduction in the number of free barbed ends available for elongation following acute Arp2/3 inhibition, thus tilting polymerization dynamics towards long linear filaments, or a compensatory shift to formin-mediated nucleation-elongation (Burke et al., 2014; Galbraith et al., 2018; Rotty et al., 2015; Spence et al., 2016). In contrast, CK-666-treated EVL KO neurons exhibited comparatively minor increases in DF length and motility (Fig. 7B,C, Video 8), demonstrating that EVL is required for the observed motility phenotypes with Arp2/3 inhibition. CK-666 also reduced F-actin content and flaring in both WT and EVL KO neurons, leaving DF as thin protrusions (Fig. 7A,B,D, Video 8). On the other hand, inhibition of formin family proteins using SMIFH2 reduced DF motility and overall dynamics in both WT and EVL KO neurons (Fig. 7A-D, Video 8). Intriguingly, following formin inhibition, retraction of small DFs and protrusions was observed in both WT and EVL KO dendrites, while Arp2/3 inhibition only reduced protrusion density in EVL KO neurons (Fig. 7A,B,E). This suggests that nascent protrusions are profoundly dependent on Arp2/3 and formin activity. Collectively, these experiments demonstrate that EVL is the primary AEF responsible for elongation of Arp2/3-dependent actin networks, and formins operate in parallel as actin nucleators and/or AEFs.

**Figure 7:**
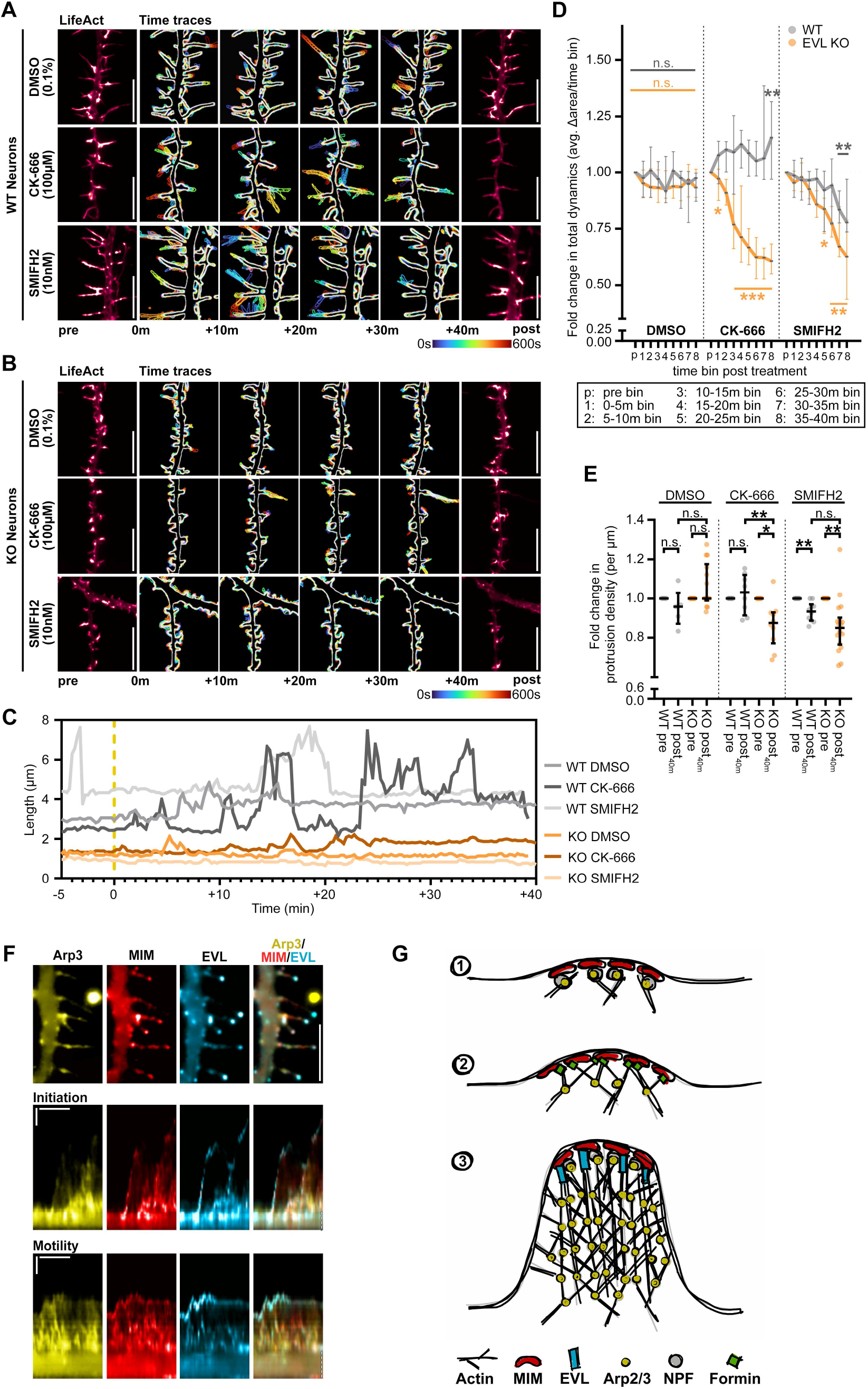
Arp2/3 and formins differentially contribute to DF initiation and motility. A-B. Segment of dendrite from WT (A) or EVL KO (B) cortical neurons at D11 expressing mRuby2-LifeAct, and treated with DMSO (0.1%), Arp2/3 inhibitor Ck666 (100µM), or pan-formin inhibitor SMIFH2 (10nM). Neurons were imaged 5 min prior to (left) and up to 40 min following (right) drug treatment. Maximum intensity projection of temporally color-coded binary mask outline of 10 min bins following treatment (middle; 15s interval, 45min duration). Scale bars = 10µm. C. Line plot of changes in length over time in a representative WT or EVL KO DF following treatment as indicated. Yellow dashed line indicates time of treatment. D. Scatterplot of WT (grey) and EVL KO (orange) total DF dynamics in 5min bins following drug treatment, and expressed as fold change relative to pre-treatment dynamics. Total dynamics were calculated as the difference in LifeAct binary mask area between timepoints, and the average of those differences within 5min time bins. E. Scatterplot of WT (grey) and EVL KO (orange) protrusion density 40min post drug treatment, and expressed as fold change relative to pre-treatment density. F. Segment of dendrite from an EVL KO cortical neuron at D11 expressing mEmerald-Arp3, MIM-mRuby2, and iRFP670-EVL (top; scale bar = 5µm), and kymographs of protein localization during DF initiation (middle) and motility (bottom). 5s interval, 2min duration, horizontal scale bar = 1min, vertical scale bar = 1µm, dashed line indicates dendrite G. Model of spatiotemporal dynamics of MIM, Arp2/3, formins, and EVL during DF initiation and protrusive motility. Statistical comparisons were made as follows: (D) two-way ANOVA and corrected for multiple comparisons. n = 7-9 neurons, N = 3 biological replicates. (E) paired Wilcoxon test, n = 9-18 dendrites, N = 3 biological replicates. * P < 0.05, ** P < 0.01, *** P < 0.001, **** P < 0.0001, n.s. is not significant. See also Supp. Video 8.

## DISCUSSION

Ena/VASP proteins are essential for neuronal morphogenesis. In this study, we identified a distinct role for EVL in the regulation of DF dynamics during early synaptogenesis in cortical neurons. EVL, through membrane-directed actin polymerization, is necessary and sufficient for mediating DF morphology and protrusive motility. In contrast, MENA is not essential for DF motility, and VASP is not highly expressed. We determined that EVL is dynamically enriched to DF tips preceding protrusion, and loss of EVL profoundly impacts DF morphogenesis and dynamics. Using an unbiased proteomics approach, we identified a MIM:EVL complex, which promotes initiation of nascent DF and their motility. As an inverse bar (I-BAR) protein, MIM facilitates membrane deformation and activates Arp2/3 to promote DF initiation (Saarikangas et al., 2015); however, the actin elongation factor(s) (AEFs) responsible for promoting actin polymerization at these initiation sites had not been identified. We provide evidence that EVL is the primary AEF responsible for elongating actin at MIM-induced proto-protrusions. Our data support the following working model of DF morphogenesis (Fig. 7F,G): (1) MIM induces an initial proto-protrusion by facilitating membrane deformation; (2) Arp2/3 is activated at proto-protrusions and, together with formin, provides the initial burst in actin remodeling leading to initiation of DF protrusion; and (3) EVL is recruited by interaction with MIM leading to rapid actin polymerization and elongation of the protrusion to form a canonical DF. In the absence of EVL, dendritic protrusions are unable to elongate, and are entirely dependent on MIM, Arp2/3, and formins. Importantly, the loss of any of these three factors in an EVL KO background causes retraction or loss of dendritic protrusions entirely.

### EVL is the major driver of DF

The critical role of Ena/VASP proteins in neuronal development is well established. MENA-VASP-EVL KO mice exhibit profound neurodevelopmental defects resulting from a failure of filopodia formation (Dent et al., 2007; Kwiatkowski et al., 2007). MENA and VASP have been shown to regulate filopodial dynamics at the growth cone during axon pathfinding (Lanier et al., 1999; Lebrand et al., 2004; McConnell et al., 2016; Menon et al., 2015; Menzies et al., 2004). In contrast, we found that neurite and axon outgrowth were unaffected by EVL KO, suggesting no growth cone defects occur with loss of EVL. On the other hand, we demonstrated that EVL is uniquely required for morphogenesis of DF. Neurons derived from EVL KO mice exhibit aberrant DF morphology and dynamics that manifest specifically during developmental periods *in vitro* when synaptic precursor DF are abundant and actively contributing to synapse formation. This indicates that DF are EVL-dependent, while earlier developmental processes are mediated by different actin regulators.

Early morphogenic processes, such as neuritogenesis and dendritic branching, are regulated by neuronal filopodia (Dent et al., 2007; Kwiatkowski et al., 2007; Portera-Cailliau et al., 2003). Prior works identified a role for VASP in the genesis of neuronal filopodia (Lin et al., 2013, 2007). Due to the examination of only early timepoints (D5-D7) in these studies, reliance upon overexpression, and the diversity in composition and function of neuronal filopodia throughout development, we are hesitant to conclude that VASP’s role in *synaptic precursor* DF is well-supported. Importantly, we found that the expression of VASP is very low in cortical neurons, compared to EVL and MENA. Interestingly, MENA KD, in contrast to EVL KD, resulted in elongated DF with higher motility, suggesting that MENA plays a role divergent from that of EVL in DF.

Furthermore, MENA’s function at the growth cone is well established (Lanier et al., 1999; Lebrand et al., 2004; McConnell et al., 2016), and in our work MENA overexpression promoted fan-like dendritic protrusions reminiscent of growth cones. We propose that MENA and VASP are important for conventional filopodia in neurons and growth cones, while EVL is uniquely responsible for DF.

### EVL recruitment and actin polymerization activity are necessary and sufficient for promoting DF motility

EVL’s function in DF, much like the DF itself, is best examined through the lens of lamellipodia biogenesis. DF are protrusions constructed of branched actin (Korobova and Svitkina, 2010), and their dynamics, like lamellipodia, are governed by Arp2/3 actin assembly and base-directed myosin-II contractility (Burnette et al., 2011; Korobova and Svitkina, 2010; Tatavarty et al., 2012). Notwithstanding the important role of Ena/VASP proteins as critical actin regulators at the leading edge and filopodia tips (Applewhite et al., 2007; Bear et al., 2002; Guercio et al., 2020), EVL’s function as a driver of membrane protrusion is not well understood. Like its sister proteins, EVL’s activity is defined by two stages: (1) recruitment, which constitutes the activation step of Ena/VASP proteins, and (2) processive actin polymerization. Recruitment, primarily via the EVH1 domain, targets EVL to polymerization sites where it binds actin and adds actin monomers, which are recruited as a profilin:G-actin complex, to the growing barbed end (Bear and Gertler, 2009). To investigate the involvement of EVL in DF, we created a panel of functional mutants that target different components of these two stages.

We demonstrated that the EVH1 domain is required for EVL’s tip localization and function in DF, where expression of ΔEVH1-EVL did not reverse the EVL KO phenotype. The CAAX-iLID system that we developed in this study allowed us to circumvent the requirement for the EVH1 domain by direct optogenetic recruitment of ΔEVH1-iLID to the membrane through photostimulation. This acute recruitment was sufficient to restore DF polymerization dynamics and length in EVL KO neurons, albeit only transiently during the time of stimulation. Furthermore, we found that deletion of EVL’s profilin binding site attenuated fast protrusion but did not entirely eliminate dynamics. Disruption of both profilin- and G-actin binding (ΔActinPoly), or actin binding entirely (ΔGAB-FAB), fully inhibited motility. Interestingly, in our hands, the GAB domain was dispensable for EVL’s function, in contrast to previous evidence that VASP-mediated actin polymerization and barbed end association requires the GAB domain (Applewhite et al., 2007; Breitsprecher et al., 2011; Hansen and Mullins, 2010). However, high concentrations of profilin:G-actin bypass GAB’s requirement (Hansen and Mullins, 2010), which may be the case in DF due to their small volume and high actin density. We have previously shown that EVL preferentially utilizes profilin-II to elongate actin filaments in breast cancer cells (Mouneimne et al., 2012). Intriguingly, profilin-II deficiency results in altered spine density and synaptic dysfunction (Ackermann and Matus, 2003; Michaelsen et al., 2010; Michaelsen-Preusse et al., 2016), phenotypes that are in line with compromised EVL function. Together, our findings support a model in which EVL is a central regulator of actin polymerization dynamics in DF.

### MIM plays a critical role in supporting EVL function in DF

I-BAR family proteins, including MIM and IRSp53, promote outward membrane curvature at sites of PIP2 enrichment (Mattila et al., 2007; Saarikangas et al., 2009), setting in motion a biophysical positive feedback loop driving PIP2 clustering and further I-BAR oligomerization (Mattila et al., 2007; McCusker, 2020; Saarikangas et al., 2009, 2015; Zhao et al., 2013). This establishes an activity hub for PIP2-responsive actin regulators to promote actin assembly and protrusion (Saarikangas et al., 2015; Senju et al., 2017). I-BAR proteins directly bind and recruit nucleation-promoting factors to activate Arp2/3-mediated actin nucleation (Lin et al., 2005; Mattila et al., 2003; Woodings et al., 2003). We have identified a tip-localized MIM:EVL complex in developing neurons that promotes DF initiation and elongation, and propose that MIM establishes hotspots of active Arp2/3 at nascent protrusions, as well as recruits and clusters EVL, to promote DF initiation and elongation.

Our findings demonstrate that MIM enrichment to foci on the dendrite licenses a proto-protrusion for elongation by EVL. During spontaneous or EVL iLID-elicited DF initiation, coincident enrichment of MIM and EVL immediately precedes protrusion, and MIM is a requisite first step in this process. When overexpressed in WT neurons, EVL predominantly elongates existing protrusions, while with MIM co-overexpression, it also substantially increases the initiation of *de novo* DF. This suggests that EVL is not sufficient for DF initiation on its own, and is limited by the availability of suitable sites along the membrane. Moreover, although MIM KD did not reduce DF dynamics, acute inhibition of MIM using MITO-iLID significantly inhibited motility compared to DF dynamics before photostimulation. This suggests that tip-localized MIM continues to promote actin polymerization in DF beyond initiation. EVL interacts with MIM via its EVH1 domain, and the two proteins are co-localized at the tips of DF during elongation. EVL KO neurons co-expressing ΔEVH1-EVL and MIM exhibited a partial restoration of motility and morphology, while expression of ΔEVH1-EVL alone was incapable of rescue. Collectively, these findings suggest that MIM creates a permissive environment for EVL’s actin polymerization activities, through the direct recruitment of EVL, and by promoting an actin environment conducive to EVL-mediated elongation.

Recent papers identified MIM as an important regulator of DF in Purkinje cells of the cerebellum and cortical neurons (Galbraith et al., 2018; Saarikangas et al., 2015). Although knockdown of MIM in EVL KO neurons strikingly reduced protrusion density in our study, it did not eliminate DF in WT neurons; this is in agreement with observations in MIM KO mouse strains (Galbraith et al., 2018; Minkeviciene et al., 2019; Saarikangas et al., 2015), suggesting possible functional redundancy with other I-BAR proteins. Indeed, IRSp53 is also implicated in dendritic spine formation, synaptic plasticity, and neurodevelopmental disorders (Choi et al., 2005; Chung et al., 2015; Kang et al., 2016). Intriguingly, Galbraith et al. provide evidence that MIM binds the formin DAAM1 and suppresses its activity in DF, while promoting activation of Arp2/3 (Galbraith et al., 2018). It is tempting to hypothesize that MIM, a multi-domain scaffolding protein, could exhibit several conformational and activity states to dynamically regulate its interaction network. Could a MIM-formin-Arp2/3 network be responsible for nucleation and DF initiation, then disruption by EVL-MIM binding promotes rapid elongation? Further studies will clarify the molecular dynamics of MIM in DF during different stages of synaptogenesis.

### EVL and formins collaborate to regulate DF motility

Formins are another class of AEF and actin nucleator implicated in DF morphogenesis. The formin mDia2 has been shown to regulate dendritic protrusion density in cortical neurons (Hotulainen et al., 2009). However, whether mDia2 is primarily acting as a nucleator-elongator, or elongates Arp2/3-nucleated filaments in DF has not been established. Acute or genetic loss of Arp2/3 results in elongated DF, likely due to a preponderance of long unbranched filaments, which have been shown to be readily suppressed by formin knockdown or inhibition (Galbraith et al., 2018; Spence et al., 2016). Formins, Ena/VASP, Arp2/3, and profilin:G-actin collaborate and compete in distinct spatiotemporal complexes to maintain actin homeostasis during cellular activities (Barzik et al., 2014; Beli et al., 2008; Rotty et al., 2015; Skruber et al., 2020; Suarez and Kovar, 2016; Suarez et al., 2015). We provide evidence that EVL and formins collaborate to regulate DF motility – EVL is required for persistent forward protrusion, but both formin and Arp2/3 activity are necessary for EVL’s function.

In EVL KO neurons, flare-like dendritic protrusions exhibit minor motility, but lacked the substantial protrusion characteristic of DF. Following acute pharmacological inhibition of formin activity, using SMIFH2, we saw a reduction in DF dynamics in WT neurons; however, in KO neurons, many dendritic protrusions withdrew entirely upon formin inhibition. This suggests that formin nucleation and/or elongation activities are required for the persistence of the protrusion. The reduction in tip motility in WT neurons with formin inhibition suggest two possible, non-exclusive models: (1) nucleation by formins and Arp2/3 occurs in the initial protrusion and DF base, and EVL regulates barbed-end elongation and protrusive motility at the tip, and/or (2) formins and EVL function as AEFs to elongate Arp2/3-nucleated filaments in parallel or under distinct regulatory mechanisms.

Furthermore, Ena/VASP proteins and active mDia2 can exist in a complex at filopodia tips in non-neuronal cells (Barzik et al., 2014); however, whether this occurs under normal conditions is unclear. Our EVL IP-MS from cultured D11 neurons did not identify any formin paralogs with high confidence, suggesting that if there is cooperation of mDia2 and EVL, they may be spatiotemporally distinct. Constitutively active mDia2, when overexpressed, localizes to DF tips (Hotulainen et al., 2009). In order to clarify the nature of these relationships, exploring the dynamics of endogenous rather than exogenously-expressed AEFs in live-cell assays is required. We have attempted to mitigate these potentially confounding issues by restoring EVL expression in a KO background, and selecting very low expressing cells for analysis. However, future works should investigate these relationships with acute manipulations of proteins at endogenous expression levels.

### Regulation of DF by EVL has potential implications for neuronal plasticity

Triple KO mice, where the expression of all three paralogs of Ena/VASP is lacking, exhibit profound defects in cortical development (Dent et al., 2007; Kwiatkowski et al., 2007). Notably, these defects were associated with compromised migration of cortical neurons, which is consistent with earlier work showing that the axon pathfinding is disrupted in MENA KO and MENA/VASP double KO mice (Lanier et al., 1999; Menzies et al., 2004). On the other hand, EVL KO mice are viable and display no gross abnormalities (Kwiatkowski et al., 2007). Together, these findings suggest that MENA and VASP, distinct from EVL, are decidedly involved in mediating large-scale cellular remodeling events, while EVL is not. Accordingly, what is EVL doing during neuronal development in the brain? The studies presented here provide evidence that among the Ena/VASP proteins, EVL play a unique role in promoting DF.

The contribution of DF to synaptogenesis in early neurodevelopment is well established (Fiala et al., 1998; Portera-Cailliau et al., 2003; Ziv and Smith, 1996), and DF continue to shape neural connectivity later in life, albeit less pervasively (Sala and Segal, 2014; Zuo et al., 2005). Experience-dependent plasticity and memory is made possible through the genesis, turnover, and structural remodeling of synapses. DF emanate from the dendrites in response to specific extracellular cues and intracellular signaling cascades (Jourdain et al., 2003; Petrak et al., 2005), and their tentative nature is thought to permit broad sampling yet refined selection of appropriate axon targets (Lohmann and Bonhoeffer, 2008; Ozcan, 2017). Synaptogenesis is frequently observed occurring via DF, however, EVL KO neurons lack DF *in vitro*, and EVL KO mice are overtly normal. What then is the evolutionary rationale for a dedicated molecular pathway that uniquely regulates DF behavior? Fine-tuning of neural connectivity requires a cadre of actin regulators to promote assembly and disassembly of synapses. Although not examined previously or in this work, it will be incredibly exciting to determine where, when, and under what circumstances EVL influences synaptic plasticity *in vivo*. Future studies will focus on understanding how EVL regulates neuroplasticity in the brain and how this affects behavior.

## Acknowledgements

We thank Seth Zimmerman (Duke University) for his technical assistance with iLID optogenetic tools, and Vasja Urbančič (University of Cambridge) for assistance with Filopodyan. We thank Jean Wilson who provided valuable feedback on the manuscript, Konrad E Zinsmaier for his guidance, and Marco Padilla-Rodriguez who provided illustrations. We thank Gillian Paine-Murrieta and the Experimental Mouse Shared Resource (EMSR) for maintenance of mouse colonies. We also thank Christopher Miller (Nikon) for his generous assistance and expertise with the microscopes. Research reported in this publication was supported by the National Cancer Institute of the National Institutes of Health under award number P30 CA023074 (to University of Arizona Cancer Center Biostatistics and Tissue Acquisition and Cellular/Molecular Analysis Shared Resources and EMSR). This work was supported by a generous fellowship from the Bisgrove Scholars Program, Science Foundation Arizona (SSP), and a National Cancer Institute grant R01 CA196885-01 (GM). Authors have no conflicts of interest to report.

## Author Contributions

Conceptualization of the project was performed by SSP and GM. Investigation was performed by SPP, ADG, PLR, and KTL. Methodology was developed by SSP, ADG, and PRL. Resources were developed by SSP. Software was developed by CWW. Project was supervised by GM, MP, PRL, and CWW. Visualization of data was performed by SSP, PRL, and GM. Writing and revision of the manuscript was performed by SSP, KTL, ADG, PRL, CW, and GM.

## Materials Availability

All unique plasmids generated in this study have been deposited to Addgene.

## Data and Code Availability

Proteomics datasets generated during this study are available at ### REPOSITORY. Uncropped western blots are available at ### MENDELEY. Custom MatLab script for DF tracking analysis available at ### GITHUB.

## Materials and Methods

### Animals

All animal procedures were performed in accordance with the regulations and protocols of the University of Arizona Institutional Animal Care and Use Committee. On embryonic day 17, pregnant mice were euthanized using CO_2_ asphyxiation. EVL knock-out mouse strain was a gift from Frank Gertler (MIT), generated as described (Kwiatkowski et al., 2007). In brief, a targeting vector disrupting exon 2 and 3 of *Evl* was electroporated into R1 embryonic stem cells for homologous recombination, and a germline clone was isolated (confirmed by Southern blot and western blot (Kwiatkowski et al., 2007)). EVL KO strains were backcrossed six times with C57B/6J to increase congenic status. Wildtype and EVL KO parents for generating embryonic primary cortical neuron cultures were derived from littermates from heterozygous crossings.

### Cell Culture

<u>For primary cortical neuron cultures</u>, the cortex was isolated from brains of embryos of either sex, dissociated, and cultured by methods described previously (Kaech and Banker, 2006). Cortical tissues were diced and digested for 20 minutes at 37°C in a solution of 0.125% Trypsin-EDTA (Corning, 25-053-Cl) and 1.0% (wt/vol) DNase (Bioline, 9003-98-9) in calcium- and magnesium-free 1X HBSS (Gibco, 14065-056). Following three five-minute rinses in HBSS, digested tissue was triturated 20 times each with three P1000 pipette tips cut to progressively smaller gauges. The homogenate was passed through a 70µm filter, counted, and cryostored as described (Parker et al., 2018). Isolated cells were resuspended in CryoStor CS10 (BioLife Solutions, 210102) to 6 million cells per mL, aliquoted, and cryostored at -80°C for at least two days then transferred to liquid nitrogen. Cryostored cells were used in all assays except mass spectrometry.

After dissection or upon thawing, primary neuronal cells were plated on standard tissue culture dishes or #1.5 coverglass-bottom dishes (MatTek, P35G-1.5-14-C) that were coated overnight with 0.001% poly-L-lysine (MilliporeSigma, P4707, diluted in water 1:10) and washed three times for 10 minutes each with water. Cells were seeded at 0.78×10^3^/mm^2^ for low-density cultures for live-imaging and immunofluorescence, and 2.60×10^3^/mm^2^ for high-density cultures for protein and RNA samples. Cells were initially plated in Plating Media, containing 5% FBS (VWR, 97068-085) and 0.6% (wt/vol) D-glucose (Fisher Scientific, BP350-500) in MEM with Earle’s salts and L-glutamine (Corning, 10-010CV). Two to four hours after plating, Plating Media was exchanged for Neuronal Maintenance Media, containing 2% NeuroCult SM1 supplement (StemCell Technologies, 05711), 1% L-glutamine (Corning, 25-005CI), and 1% penicillin-streptomycin (Corning, 30-002CI) in Neurobasal Media (Thermo Fisher Scientific, 21103049). Primary neuron cultures are maintained at 37°C and 5% CO_2_. One-half volume media exchange was given on day four (D4), and one-third media exchanges every three to four days thereafter, using Neuronal Maintenance Media pre-conditioned on glial cells maintained in a separate culture, and supplemented with 2% NeuroCult and 5µM cytosine arabinoside (AraC, MilliporeSigma, C6645) to curb glial cell proliferation. One-third media exchange was given prior to imaging, and all experiments were performed on D11 unless indicated otherwise.

<u>For lentivirus production</u>, HEK293T were cultured in 10% FBS (VWR, 97068-085), 1% penicillin-streptomycin (Corning, 30-002CI), and 1% L-glutamine (Corning, 25-005CI) in high glucose DMEM with sodium pyruvate (Corning, 10-013-CV). Plastic cell culture dishes were coated with poly-D-lysine (MilliporeSigma, P0899, 0.1 mg/ml)) to promote cell adhesion during lentiviral production. Cells are maintained at 37°C and 5% CO_2_. 60% confluent HEK293T cells were transfected using PEI (Polysciences, 23966) in OptiMEM (Thermo, 31985070) as previously described (Yang et al., 2017) with transfer plasmid, pMD2.G, and psPAX2 (gifts from Didier Trono, Addgene #12259 and #12260; RRID:Addgene_12259 and RRID:Addgene_12260) at a 1:0.25:0.75µg plasmid ratio and 3:1 PEI:DNA ratio. Viral supernatant was collected 48-72 hours post-transfection, filtered through 0.45µm filters, and added directly to primary neuron cultures. All lentiviruses were added four days before experiments with the exception of LifeAct viruses which were added at the time of plating. No antibiotic selection was used in transduced neuronal cultures.

<u>Pharmacological Agents</u> to induce expression of pCW57.1 mCherry-FP4/AP4-MITO, 1µg/mL doxycycline (MilliporeSigma, D9891) was added to culture media 12 hours before imaging experiments. Arp2/3 nucleation inhibitor Ck-666 (100µM in DMSO, MilliporeSigma, SML0006) or formin inhibitor SMIFH2 (10µM in DMSO, MilliporeSigma, 344092) was acutely added during live-imaging, or cells were pre-treated for 30 minutes prior to inhibitor-optogenetic experiments.

### Plasmids and Cloning

#### Sources

NEB Stables (New England BioLabs, C3040H) were used to propagate all lentiviral transfer plasmids to reduce recombination. NEB5α (NEB, C2987H) were used to propagate any non-lentiviral expression plasmids. Plasmids containing source cDNA sequences for *Mus musculus* EVL, MENA, and ARP3 were gifts from Frank Gertler (MIT, (Carl et al., 1999)). Plasmid containing source cDNA sequence for *Mus musculus* MIM (NM_144800) with C-terminal myc- and FLAG-tag was purchased from Origene (MR210506). All plasmids were sequenced to confirm the correct coding sequence (Eton Bioscience).

#### Cloning

All-in-one <u>doxycycline-inducible lentiviral transfer plasmids</u> (pLV-Dox mCherry-FP4/AP4-MITO) were generated by PCR and subcloning mCherry-FP4-MITO or mCherry-AP4-MITO (a gift from James Bear, UNC, (Bear et al., 2000)) in place of Cas9 in pCW-Cas9 (a gift from Eric Lander & David Sabatini, (Wang et al., 2014), Addgene #50661; RRID:Addgene_50661). Briefly, Cas9 was cut out and replaced with a multiple cloning site by annealed oligo cloning, and AgeI-BamHI was used to insert FP4 sequences. <u>TurboRFP-expressing pLKO transfer plasmid</u> was generated by PCR and subcloning TurboRFP in place of the puromycin resistance gene at BamHI-KpnI in pLKO.1 - TRC cloning vector (a gift from David Root, (Moffat et al., 2006), Addgene #10878; RRID:Addgene_10878). Validated *Mus musculus* <u>shRNA sequences</u> were obtained from The RNAi Consortium (The Broad Institute via MilliporeSigma, (Moffat et al., 2006)) and oligos were ligated between the AgeI-EcoRI sites (replacing the 1.9kb stuffer) of our pLKO.1 TurboRFP cloning vector using annealed oligo cloning. shRNA sequences and source MilliporeSigma product number are given in Key Resources Table. <u>Lentiviral LifeAct expression vectors</u> were published previously (Padilla-Rodriguez et al., 2018; Parker et al., 2018): pLenti LifeAct-EGFP BlastR (Addgene #84383; RRID:Addgene_84383), pLenti-LifeAct-mRuby2 BlastR (Addgene #84384; RRID:Addgene_84384), pLenti LifeAct-iRFP670 BlastR (Addgene #84385; RRID:Addgene_84385). <u>Lentiviral cDNA expression vectors</u> were generated by PCR and subcloning cDNA of interest into the transfer plasmids pLenti CMVie-IRES-BlastR or pLenti CMVie-IRES-BlastR alt MCS (pCIB) published previously (Puleo et al., 2019) (Addgene #119863 and #120862; RRID:Addgene_119863 and RRID:Addgene_120862). To reduce expression of EVL, the CMVie promoter was replaced with the Ef1a short promoter EFS in some constructs. cDNAs were tagged with mEmerald (a gift from Michael Davidson, Addgene #53975; RRID:Addgene_53975), mRuby2 (a gift from Michael Davidson, (Lam et al., 2012), Addgene #54768; RRID:Addgene_54768), or piRFP670 (a gift from Vladislav Verkhusha, (Shcherbakova and Verkhusha, 2013), Addgene #45457; RRID:Addgene_45457) on the N-terminus of *EVL* or *Arp3*, or C-terminus of *Mtss1*. For EVL iLID and MIM iLID optogenetic systems, tgRFPt-SspB(R73Q) was PCRed and subcloned from pLL7.0 mTiam1(64-437)-tgRFPt-SSPB R73Q (a gift from Brian Kuhlman, (Guntas et al., 2015), Addgene #60418; RRID:Addgene_60418) and added to the N-terminus of *Evl* or C-terminus of *Mtss1*. Lentiviral expression vectors for MIM-mRuby2, MIM-iRFP670, and MIM iLID were modified to remove the IRES-blasticidin resistance cassette to reduce plasmid size. Domain deletion mutants of EVL were generated by PCR or inverse PCR and self-ligation cloning. All plasmids created for this paper will be made available through Addgene. Complete list of plasmids used in this paper can be found in the Key Resources Table.

### mRNA Extraction and RT-qPCR

Total RNA content was isolated from primary neuron cultures using Trizol (Life Technologies, 15596026) and Direct-zol RNA MiniPrep kit (Zymo Research, R2050), according to the manufacturer’s instructions, including the optional DNase digestion step to eliminate genomic DNA in the sample. cDNA was synthesized using 1000ng of input RNA and qScript cDNA Supermix (Quantabio, 84033) according to the manufacturer’s instructions. RT-qPCR reactions were run in triplicate using an ABI 7500 Fast Real-Time PCR System (Applied Biosystems) and Apex qPCR 2X Master Mix Green, Low ROX (Apex BioResearch Products, 42-119PG). Primer pairs were confirmed to have 85-110% efficiency, which was determined from the slope of the best fit curve for the C_T_ of cDNA serial dilutions. C_T_ values were normalized to the average of three control genes: *Gapdh, Eef1a1*, and *Rpl29*. Relative copy numbers were determined using comparative CT method (2^-ΔCT^; (Schmittgen and Livak, 2008)). Primer sequences are given in Key Resources Table.

### Western Blotting

Cells were lysed with ice cold lysis buffer buffer (25mM HEPES pH 7.4, 150mM NaCl, 1% NP-40, 0.25% Na Deoxycholate, 10% glycerol) supplemented with protease and phosphatase inhibitor cocktails (MilliporeSigma, 539134; Boston BioProducts, BP-479), and incubated on ice for five minutes. Cellular debris was pelleted out by centrifugation at 21,000g for 15 minutes at 4°C. Protein concentration was determined using a Bradford assay compared to BSA protein standards. Supernatant was mixed with Laemmli buffer and boiled at 95°C for five minutes. To determine protein expression, 20-30µg of protein lysate was loaded per well. For immunoprecipitation blots, 5% of input lysate and entire IP eluate was loaded. Samples were resolved using SDS-PAGE and 10% acrylamide gels. Resolved samples were transferred to nitrocellulose membranes (LI-COR, 926-31090), and membranes were blocked in Intercept Blocking Buffer (LI-COR, 927-70001) for one hour rocking at room temperature. Membranes were probed overnight at 4°C with primary antibodies diluted in blocking buffer and 0.2% Tween-20. After washing in TBS with 0.1% Tween-20, membranes were probed with near-infrared fluorescent secondary antibodies diluted in blocking buffer and 0.2% Tween-20 for one hour at room temperature. Antibodies were as follows: rabbit polyclonal anti-EVL (a gift from Frank Gertler, 1:1000), rabbit polyclonal anti-ENAH (Sigma Prestige HPA028696, 1:250), rabbit monoclonal anti-VASP (Cell Signaling #3132, 1:500), rabbit polyclonal anti-MTSS1 (Thermo, PA5-23200, 1:500) rabbit polyclonal anti-FLAG (ProteinTech Group, 20543-1, 1:1000), rabbit polyclonal anti-GFP (ProteinTech Group, 66002-1 1:1000), mouse monoclonal anti-actin (ProteinTech Group, 66009-1, 1:5000), goat anti-mouse Alexa Fluor 680 and goat anti-rabbit Alexa Fluor 790 secondary antibodies (Thermo, A21057 and A11367 1:20,000).

### Immunoprecipitation and Mass Spectrometry

<u>For immunoprecipitation-western blot experiments</u>, 10cm plates of 60% confluent HEK293T were transfected with 4µg of each plasmid (total of 8µg), and 24µg PEI. 24 hours post transfection, cells were lysed with ice cold IP buffer (10% glycerol, 1% NP-40, 50mM Tris pH 7.5, 200mM NaCl, 2mM MgCl_2_) supplemented with protease and phosphatase inhibitor cocktails (MilliporeSigma, 539134; Boston BioProducts, BP-479) and incubated on ice for five minutes. Cellular debris was pelleted out by centrifugation at 21,000g for 15 minutes at 4°C. Protein concentration was determined using a Bradford assay compared to BSA protein standards, and 1-2mg of protein was loaded into each IP reaction. Lysates were pre-cleared in 5µL of magnetic Protein A/G bead slurry (Thermo, 88802), and 2.5% of the volume of whole cell extract was set aside. Reactions were incubated overnight at 4°C with gentle rotation with mouse monoclonal anti-FLAG (ProteinTech Group, 66008-3, 5µg antibody per mg protein). 25µL magnetic Protein A+G bead slurry was added to each reaction, and incubated for two hours at 4°C with gentle rotation. Beads were washed three times in IP buffer with 0.1% Tween-20, and proteins were eluted from beads by boiling at 95°C for ten minutes in 2X Laemmli buffer diluted in IP buffer. Samples were resolved by SDS-PAGE and western blotting as described above.

<u>For affinity purification-mass spectrometry experiments</u>, two poly-L-lysine-coated 15cm plates were plated with ten million primary neuron cells. Cells were lysed on D11 and protein samples were handled as described above. Reactions were incubated overnight at 4°C with gentle rotation with mouse control IgG (Santa Cruz, sc-2025) or mouse monoclonal anti-EVL (a gift from Frank Gertler) at 5µg antibody per mg protein. 25µL magnetic Protein A/G bead slurry (Thermo, 88802) was added to each reaction, and incubated for two hours at 4°C with gentle rotation. Beads were washed three times in IP buffer, and proteins were eluted from beads by boiling at 95°C for ten minutes in 2X laemmli buffer diluted in IP buffer. Entire eluate was loaded onto Bolt 4 to 12% Bis-Tris pre-cast gels (Thermo, NW04120BOX), resolved by SDS-PAGE, and the gels were stained with Bio-Safe Coomassie G-250 Stain (Bio-Rad, #1610786; Hercules, CA).

#### In-gel digestion

in-gel tryptic digestion was performed as previously described (Kruse et al., 2017; Parker et al., 2019). In brief, each lane of the SDS-PAGE gel was cut into eight slices. Each gel slice was placed in a 0.6 mL LoBind polypropylene tube (Eppendorf), destained twice with 375 µL of 50% acetonitrile (ACN) in 40mM NH_4_HCO_3_ and dehydrated with 100% ACN for 15 min. After removal of the ACN by aspiration, the gel pieces were dried in a vacuum centrifuge at 60°C for 30 minutes. Trypsin (250 ng; Sigma-Aldrich) in 20 μL of 40mM NH_4_HCO_3_ was added, and the samples were maintained at 4°C for 15 minutes prior to the addition of 50-100 μL of 40 mM NH_4_HCO_3_. The digestion was allowed to proceed at 37°C overnight followed by termination with 10 μL of 5% formic acid (FA). After further incubation at 37°C for 30 minutes and centrifugation for 1 minute, each supernatant was transferred to a clean LoBind polypropylene tube. The extraction procedure was repeated using 40 μL of 0.5% FA, and the two extracts were combined and dried down to ∼5-10 μL followed by the addition of 10 μL 0.05% heptafluorobutyric acid/5% FA (vol/vol) and incubation at room temperature for 15 minutes. The resulting peptide mixtures were loaded on a solid phase C^18^ ZipTip (Millipore, Billerica, MA) and washed with 35 μL 0.005% heptafluorobutyric acid/5% FA (vol/vol) followed by elution first with 4 μL of 50% ACN/1% FA (vol/vol) and then a more stringent elution with 4 μL of 80% ACN/1% FA (vol/vol). The eluates were combined and dried completely by vacuum centrifugation and 6 μL of 0.1% FA (vol/vol) was added followed by sonication for 2 minutes. 2.5 μL of the final sample was then analyzed by mass spectrometry.

#### Mass spectrometry and database search

HPLC-ESI-MS/MS was performed in positive ion mode on a Thermo Scientific Orbitrap Fusion Lumos tribrid mass spectrometer fitted with an EASY-Spray Source as previously described (Parker et al., 2019). NanoLC was performed using a Thermo Scientific UltiMate 3000 RSLCnano System with an EASY Spray C^18^ LC column (Thermo, 50 cm × 75 μm inner diameter, packed with PepMap RSLC C18 material, 2 μm, cat. # ES803); loading phase for 15 minutes; mobile phase, linear gradient of 1–47% ACN in 0.1% FA for 106 minutes, followed by a step to 95% ACN in 0.1% FA over 5 minutes, hold 10 minutes, and then a step to 1% ACN in 0.1% FA over 1 minute and a final hold for 19 minutes (total run 156 minutes); Buffer A = 100% H_2_O in 0.1% FA; Buffer B = 80% ACN in 0.1% FA; flow rate, 250-300 nL/min. All solvents were liquid chromatography mass spectrometry grade. Spectra were acquired using XCalibur, version 2.1.0 (Thermo). A “top 15” data-dependent MS/MS analysis was performed (acquisition of a full scan spectrum followed by collision-induced dissociation mass spectra of the 15 most abundant ions in the survey scan). Dynamic exclusion was enabled with a repeat count of 1, a repeat duration of 30 seconds, an exclusion list size of 500, and an exclusion duration of 40 seconds. Tandem mass spectra were extracted from Xcalibur ‘RAW’ files and charge states were assigned using the ProteoWizard 2.1.x msConvert script using the default parameters. The fragment mass spectra were searched against the 2016 *Mus musculus* SwissProt database (16838 entries) using Mascot (Matrix Science, London, UK; version 2.4) using the default probability cut-off score. The search variables that were used were: 10 ppm mass tolerance for precursor ion masses and 0.5 Da for product ion masses; digestion with trypsin; a maximum of two missed tryptic cleavages; variable modifications of oxidation of methionine and phosphorylation of serine, threonine, and tyrosine. Cross-correlation of Mascot search results with X! Tandem was accomplished with Scaffold (version Scaffold_4.8.7; Proteome Software). Probability assessment of peptide assignments and protein identifications were made using Scaffold. Only peptides with ≥ 95% probability were considered.

### Single Cell RNA-seq

Mouse and human brain single cell RNA-seq data was downloaded from the Allen Brain Map data portal (© 2010 Allen Institute for Brain Science. Allen Brain Map. (Sunkin et al., 2013) Available from: https://portal.brain-map.org/atlases-and-data/rnaseq). In our analyses, we used read counts derived from the 2019 Smart-Seq method. The 2019 Smart-Seq data was selected due to having an overall greater read depth than the 2020 10x Genomics data. To directly compare the gene expression of *Evl, Enah*, and *Vasp* within mouse glutamatergic neurons, we first selected the top 1,000 glutamatergic neurons that had the greatest number of read counts. Next, we performed TPM normalization on the expression matrix using the R function “calculateTPM” from the R library “scater” (McCarthy et al., 2017). The gene lengths used in the calculateTPM function were calculated from the GTF2LengthGC.R script provided by the github user dpryan79. For the GTF and FASTA input files, gencode.vM25.annotation.gtf and GRCm38.p6.genome.fa files were downloaded from GENCODE (Frankish et al., 2018). The same pipeline was used to calculate gene expression of *EVL, ENAH* and *VASP* in human glutamatergic cells, except the GENCODE files gencode.v34.annotation.gtf and GRCh38.p13.genome.fa were used to calculate gene lengths.

To compare the gene expression of *Evl, Enah*, and *Vasp* between glutamatergic neurons and non-neuronal cells in mice, we first selected the top 250 glutamatergic neurons and the top 250 non-neuronal cells that had the highest read counts. Next, the SCnorm pipeline was used to normalize gene expression across samples (Bacher et al., 2017). All default settings were used except the parameter “ditherCounts” was set to TRUE. The same pipeline was utilized for the comparison of *EVL, ENAH* and *VASP* between human glutamatergic neurons and non-neuronal cells.

### Immunofluorescence

Neurons were seeded on poly-L-lysine-coated #1.5 coverslips (Carolina Biological Supply, 633029). At indicated timepoints, coverslips were briefly washed in Dulbecco’s PBS with calcium and magnesium (Caisson Labs, PBL02), and fixed for 15 minutes with 4% paraformaldehyde (diluted in PBS from 16%, Electron Microscopy Services, 15710) and 4% sucrose in PBS at 37°C. Subsequent steps were performed at room temperature. Autofluorescence was quenched by incubation with 50mM NH_4_Cl for 10 minutes at room temperature. Coverslips were blocked and permeabilized using a buffer of 10% goat serum and 0.1% Triton X-100 in PBS and incubating for 30 minutes. Antibodies were diluted in a buffer of 3% goat serum and 0.1% Triton X-100 in PBS. Coverslips were incubated with primary antibodies for one hour, washed three times in PBS with 0.1% Tween-20, and incubated with secondary antibodies and Alexa Fluor 488 phalloidin (Thermo, A12379, 1:40) for 30 minutes. Coverslips were mounted using Prolong Gold Antifade (Thermo, P36930), and allowed to cure for at least 24 hours prior to imaging. Antibodies used were as follows: rabbit monoclonal anti-βIII-tubulin (Cell Signaling #5568, Clone D71G9; 1:100), rabbit monoclonal anti-MAP2 (Cell Signaling #8707, Clone D5G1; 1:500), goat anti-rabbit Alexa Fluor 568 (Thermo, A11011, 1:1000), goat anti-rabbit Alexa Fluor 647 (Thermo, A21245, 1:1000).

### Microscopy

All images were collected on a Nikon Ti-E Total Internal Reflection Fluorescence (TIRF) microscope (Nikon Instruments) equipped with an ORCA-Flash 4.0 V2 sCMOS camera (Hamamatsu), motorized stage, perfect focus system, and environmental chamber (InVivo Scientific) to maintain humidity, 37°C and 5% CO2 ambient conditions. <u>For widefield fluorescence imaging</u>, microscope is equipped with a SOLA solid-state LED white light source (Lumencor, 100% power), and a DAPI/FITC/CY3/CY5 excitation, emission, and dichroic filter set (89000 Sedat Quad ET, Chroma. Excitation Filters: D350/50x, ET402/15x, ET490/20x, ET555/25x, ET645/30x. Emission Filters: ET455/50m, ET525/36m, ET605/52m, ET705/72m). <u>For TIRF imaging</u>, microscope is equipped with a 405/488/561/640nm laser launch (Nikon, LU-n4, 15mW, 100% power except for optogenetic activation at 10% power), Ti-TIRF-E Motorized Illuminator Unit, and utilized with C-FL TIRF Ultra Hi S/N 405/488/561/638 Quad Cube, Z Quad HC Cleanup, and HC TIRF Quad Dichroic. Acquisition software was NIS Elements (Nikon).

#### Live-cell imaging

Primary neuron cultures were plated on poly-L-lysine-coated coverglass bottom dishes as described under *Cell Culture*. Neurons were imaged using TIRF microscope, which we determined to be the optimal method available to us to minimize phototoxicity, improve resolution and maximize acquisition speed. Images were acquired using a 100x Plan Apo TIRF 1.49NA objective (Nikon). Images were acquired on a five second interval for five minutes, except for pharmacological treatments in which three positions were acquired on a 15 second interval for 45 minutes, and optogenetic activation experiments which were acquired on a five second interval for 15 minutes. Images were acquired with 1×1 binning, or 2×2 binning for optogenetic activation experiments, and wherever more than two fluorescent channels were acquired (MIM+EVL+LifeAct experiments, MIM+EVL+ARP3 experiments) in order to decrease the light dose. Exposure times ranged between 70ms and 200ms at 100% laser power at TIR critical angle for each channel. <u>For optogenetic activation experiments</u>, whole cells were illuminated with vertical incident 488nm laser light (non-TIRF) at 10% power for 400ms exposure every five seconds during the stimulation phase.

#### Fixed cells

Primary neuron cultures were plated on poly-L-lysine-coated coverslips as described under *Cell Culture* and *Immunofluorescence*. Cells were imaged using 40X Plan Fluor 1.3NA objective (Nikon). 5×5 fields of images were acquired and stitched to increase field of view for unbiased morphological analysis.

### Software and Image Analysis

#### Processing

Time-lapse image stacks were processed using NIS Elements AR (Nikon). Pre-processing steps for analysis included the following workflow: correcting for drift in time using “Align ND Document”, correcting for photobleaching using “Equalize Intensity in Time” to the first frame’s histogram, and segmenting the neuron by merging denoised duplicates of fluorescent channels into a single binary mask, and subtracting the background. This eliminated any non-cell background from mean tip intensity measurements, which were measured from a 384nm radius about the tracked tip. All fluorescence intensity analysis was performed on images without further processing. Image processing for figure preparation included the following additional workflow to improve signal to noise and presentation of live-cell imaging datasets: low-pass filtering (keep details larger than 2 pixels), either Noise2Void (Fig. 2-4, using ZeroCostDL4Mic (Chamier et al., 2021; Krull et al., 2019)) or “Advanced Denoising” (Fig. 5-7, NIS Elements), 2D Blind Deconvolution (1 iterations, low noise, calculated PSF, NIS Elements) on LifeAct channel, and 2D Richardson-Lucy Deconvolution (2 iterations, low noise, calculated PSF, NIS Elements) for all other fluorescent channels. Figure 2 was additionally processed with Rolling Ball background subtraction (1µm radius). Advanced Denoising settings were adjusted as appropriate individually due to variation in construct expression levels.

#### Analysis

<u>Tip tracking</u> was performed using Manual Tracking in NIH ImageJ (FIJI build, (Schindelin et al., 2012)). DF were only tracked if they met the following conditions: no contact with axons, neighboring DF, or debris during the time course; emanated from dendrites at least 50µm away from the center of the soma; clearly visible by brightfield during time course; if they initiated or retracted during imaging, non-existent timepoints were removed from further analysis; buckling and wagging DF were included in tracking. Using the manually tracked positions of the DF base and tip, the image files were then further analyzed with a custom MATLab script to determine the centerline path along each DF. This script used the fluorescent intensity in either the LifeAct or GFP space-filler channel in the vicinity of the tip and base coordinates to define the average tangent direction of the long axis of the DF by computing the tangent angle *q* at pixel *i* using

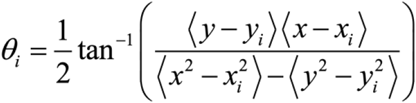

where brackets denotes the intensity-weighted average over a 15×15 pixel domain centered on the *i*th pixel. The centerline curve (*x*(*s*),*y*(*s*)) was then determined by solving

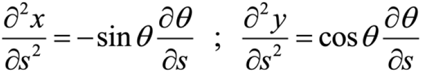

subject to the constraint that the starting and ending positions were the tracked positions of the base and tip of the DF. Using the centerline curves for each DF at each time point, we then calculated the absolute tip displacement, DF length, and mean tip fluorescence intensity and were able to extract the following metrics: <u>average filopodial tip speed</u> calculated as the average of the instantaneous speeds (absolute tip displacement per 5s interval) between successive timepoints; <u>percent motile</u>, percent of total DF population with average tip speeds greater than 0.0128µm/s (motile) or less than 0.0128µm/s (non-motile); <u>percent time motile</u>, the percent of time per DF in which instantaneous speed was greater than 0.0128µm/s; <u>average length</u>, the distance from base to tip along the centerline curve, <u>median protrusion or retraction rate</u>, the positive or negative change in length between successive timepoints, when instantaneous change in length was greater than +/-0.0128µm/s (motile); <u>mean fluorescence intensity</u> for a circular area of 384nm radius surrounding the distal DF tip; <u>fluorescence intensity variance</u>, a measure of the spread of intensity values compared to the mean. Fluorescence intensity values were normalized for expression by the minimum local intensity during the duration of imaging. For defining motile versus non-motile filopodia, or substantiative protrusion/retraction rates, a threshold of 0.0128µm/s was chosen as it represents one pixel (effective size at 100X=0.064µm) displacement per five second interval and undistinguishable from tracking error. <u>Neurite morphology</u> was measured using the ImageJ plug-in Simple Neurite Tracer (Longair et al., 2011). Tracings were used to determine the number and length of primary and higher order neurites, and length of the axon (the longest Tau-positive process). <u>Protrusion density</u> was determined by counting proturbences along a length of dendrite. <u>Overall dynamics</u> were determined by generating a binary mask of LifeAct signal for segments of dendrite, averaging the binary areas within indicated time bins, and examining the fold change in area following pharmacological treatment.

### Statistical Analysis

#### Filopodyan

We utilized the R Studio package of Filopodyan (Urbančič et al., 2017), a filopodia tracking package to determine the correlation of MENA or EVL tip fluorescence intensity to various DF metrics and the cross-correlation function and offset. Data output from our custom MatLab script was reformatted for compatibility with Filopodyan inputs. R Studio scripts were edited for pixel size, interval, and to permit compatibility and data export. Filopodyan_Masterscript.R comprising Modules 1, 2, and 3 was used to calculate and configure input data for FilopodyanR_CCF.R, CCF_subcluster-analysis.R, and CCF_Randomisations.R. FilopodyanR_CCF.R calculates the cross-correlation function (CCF) of Tip Fluorescence (FTIP; mean fluorescence intensity at tip region) and Direction-Corrected Tip Movement (DCTM; protrusion and retraction as calculated by changing length), calculated from a moving average of three adjacent timepoints. Using unbiased hierarchical clustering, high CCF DF clusters and low CCF DF clusters were designated as the top-correlating subcluster (TCS) and non-TCS, respectively. CCF_subcluster-analysis.R was used to extract and analyze DF metrics for the TCS and non-TCS populations. CCF_Randomisations.R generated 1000 randomized datasets by shuffling eight-timepoint-blocks of motility data with respect to tip fluorescence data for each DF, and performing identical cross-correlation analysis on datasets with similar subcluster size. Specific statistical analyses are given in the Figure Legends.

#### General

Statistical analyses were performed using Prism v9.1.0 (GraphPad). All data were tested for normality using the D’Agostino-Pearson normality test. Outliers were not removed for any statistical comparisons. Unpaired parametric comparisons were made using the Student’s t-test or Welch’s t-test. Unpaired non-parametric comparisons were made using the Mann-Whitney test or Kruskal-Wallis test. Paired data, including drug treatment and optogenetic stimulation data, were non-parametric and analyzed by Friedman’s Test or Wilcoxon Test. Where appropriate, analyses were corrected for multiple comparisons by Dunn’s correction.

## Figure Legends

**Supplemental Figure 1:**
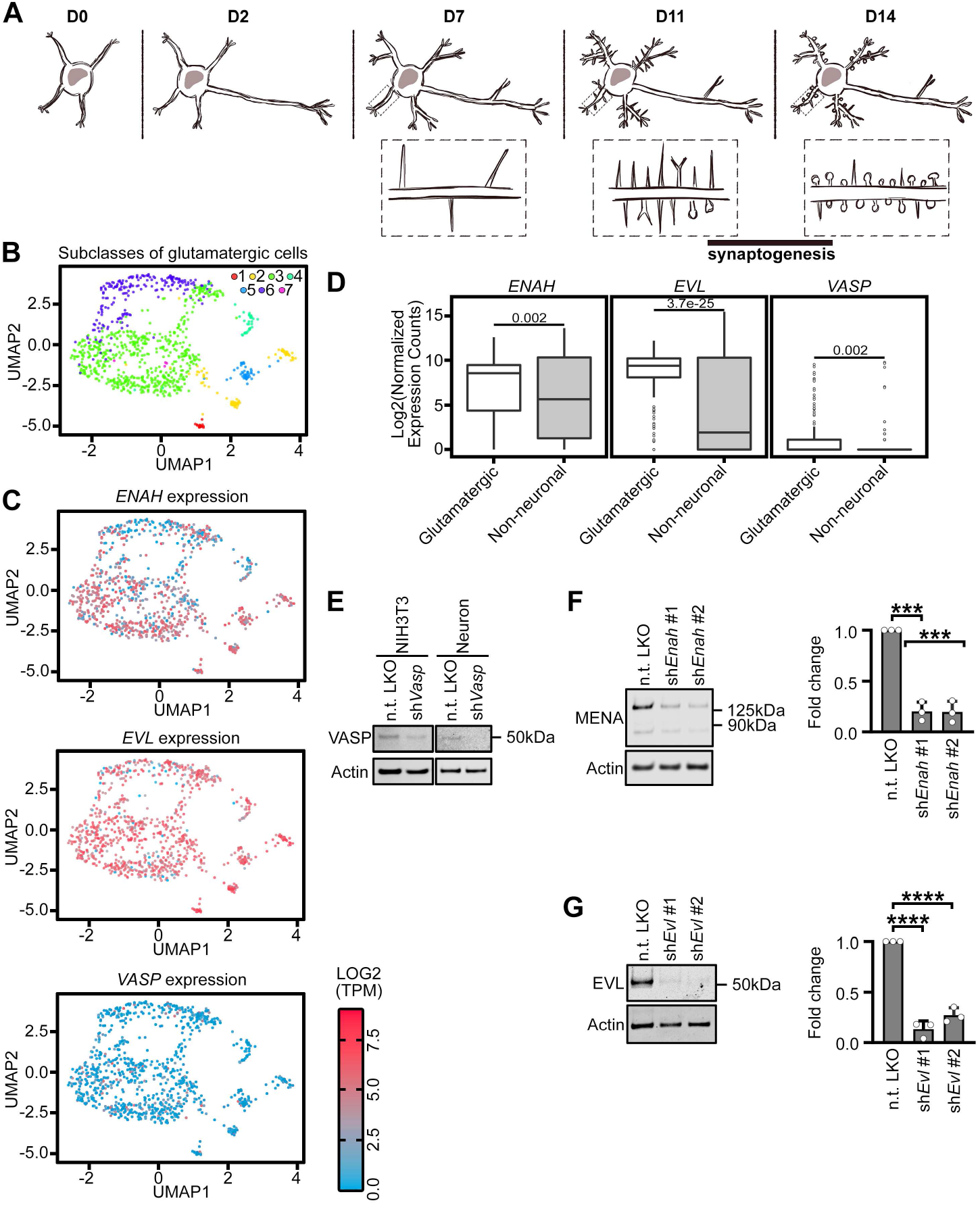
EVL is the predominant Ena/VASP paralog regulating dendritic filopodia. A. Schematic of *in vitro* cortical neuron development. B. Uniform manifold approximation and projection (UMAP) subcluster categories of single-cell RNAseq expression profiles from human glutamatergic neurons. 1: L5/6 near-projecting cortex (CTX), 2: L6 corticothalamic CTX, 3: intratelencephalic (IT) CTX, 4: L5/6 IT CTX *Car3*, 5: L6b CTX, 6: L4 IT CTX, 7: L5 IT CTX. C. Log2(TPM) expression in human of *ENAH, EVL*, and *VASP* across single-cell RNAseq UMAP subcluster categories. D. Comparison of *ENAH, EVL*, and *VASP* expression in human across single-cell RNAseq grouped by glutamatergic versus non-neuronal cell types. Statistical comparisons of RNA abundance in cell type categories made by Welch’s t-test, n = 250 cells. E. Representative western blot confirming specificity of VASP antibody in NIH3T3 (left) and primary neuron cultures (right) with pLKO-shRNA-TurboRFP targeting *Vasp* mRNA. F,G Representative western blots of protein lysates from primary neurons at D11, transduced on D7 with indicated pLKO-shRNA-TurboRFP lentiviral particles targeting *Enah* (F) or *Evl* mRNA (G; left). Fold change quantification of knockdown compared to non-targeting shRNA control (right). Statistical comparisons of shRNA knockdown efficiency were made by unpaired t-test. N = 3 biological replicates. * P < 0.05, ** P < 0.01, *** P < 0.001, **** P < 0.0001, n.s. is not significant. See also Fig. 1, Supp. Video 1.

**Supplemental Figure 2:**
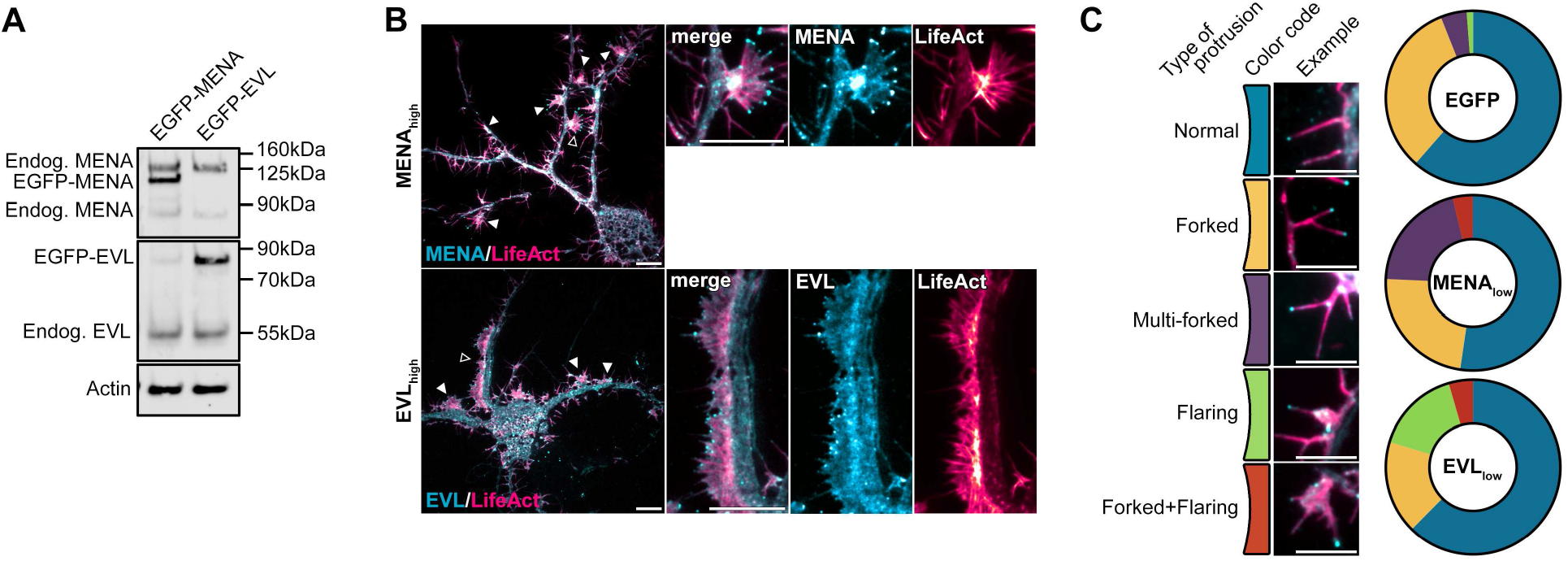
MENA and EVL overexpression enhance DF motility. A. Representative western blot of protein lysates from D11 mouse cortical neuron cultures expressing EGFP-MENA or EGFP-EVL, and probed with antibodies targeting MENA or EVL for detection of endogenous and overexpressed species. B. Examples of extreme phenotypes observed in live primary mouse cortical neurons at day *in vitro* 11 (D11) expressing mRuby2-LifeAct with high expression (SNR>1.5) of EGFP-MENA or EGFP-EVL. Left: full cell image. Closed arrowheads indicate abnormal structures. Open arrowhead indicates magnified region (right). C. Examples and quantification of DF morphological phenotypes observed in D11 neurons expressing mRuby2-LifeAct with low-expression (SNR<1.5) of EGFP-MENA or EGFP-EVL. N = 83-133 DF per condition, N = 3 neurons. Scale bars = 10µm. See also Fig. 2.

**Supplemental Figure 3:**
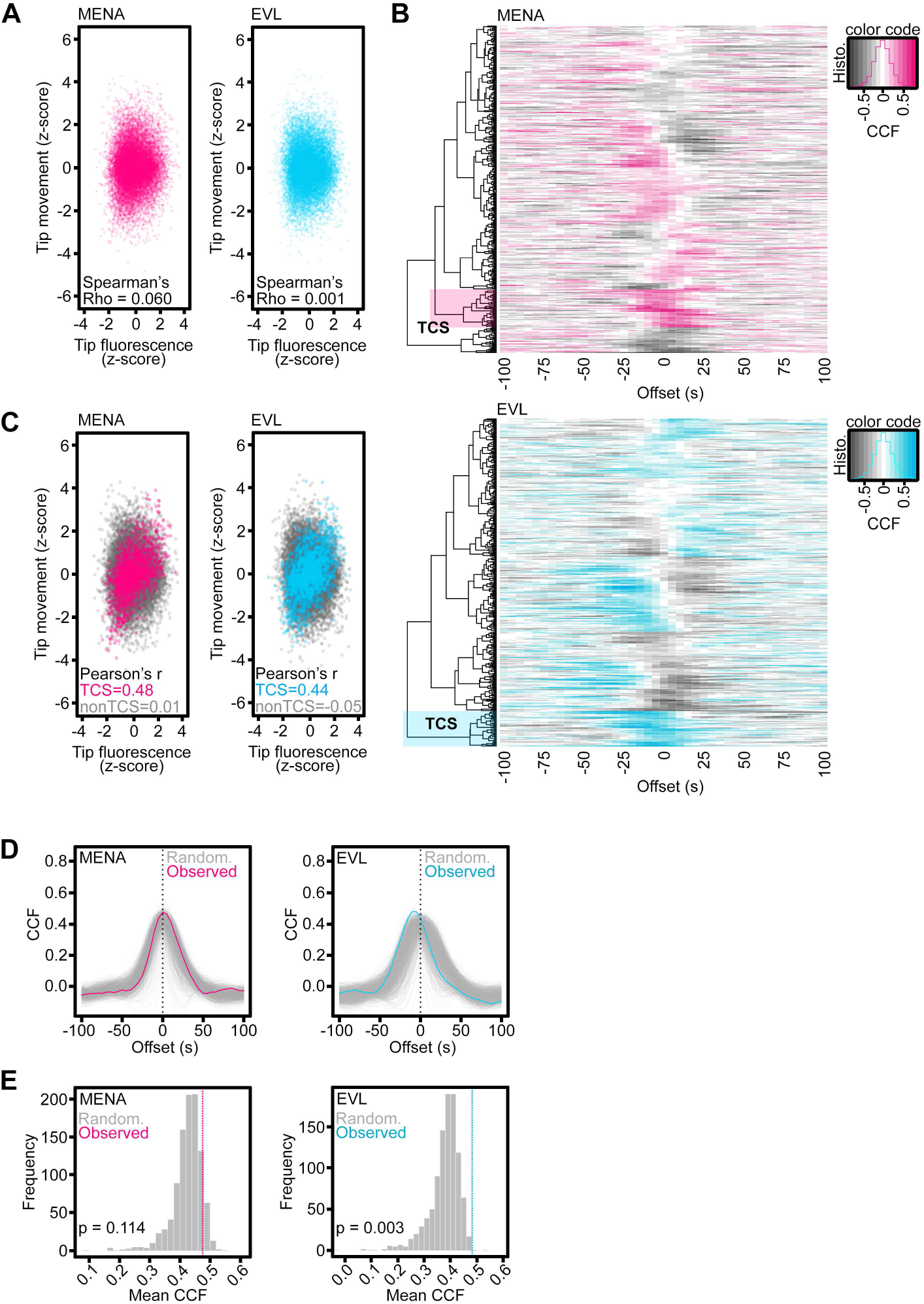
Tip enrichment of EVL precedes DF protrusion. A. Scatterplots of tip fluorescence and tip motility z-scores at individual timepoints across entire population of analyzed DFs from D11 neurons expressing EGFP-MENA or EGFP-EVL. Poor correlation of tip fluorescence and motility observed by Spearman’s correlation test across total population of DFs. B. Heatmaps of cross-correlation function (CCF) of normalized tip fluorescence intensity and tip motility per individual DF (rows) as a function of time offset (columns), of DFs from D11 neurons of indicated conditions. Positive CCFs indicated in color, negative CCFs indicated in grey. High CCFs occurring with negative offset values indicates that fluorescence enrichment precedes motility; high CCFs with positive offset values indicates fluorescence enrichment follows motility. Color shaded region denotes top-correlating subcluster (TCS) of DFs within each condition. This subpopulation was used to compare metrics for DFs in which fluorescence intensity and motility are highly-correlated (TCS), versus DFs in which this correlation is weak or negatively correlated (non-TCS). Histogram (overlaid on color code scale bar) displays DF population distribution across CCF values. C. Scatterplots of tip fluorescence and tip motility z-scores at individual timepoints for TCS (color) overlaid on non-TCS (grey) DFs of indicated conditions. Strong correlation observed by Pearson’s correlation test for TCS compared to non-TCS of DFs. D. Line plots of CCF of normalized tip fluorescence intensity and tip motility as a function of time offset for observed TCS (color lines) and the TCS from randomized datasets (grey lines). Randomization was performed by shuffling eight-timepoint blocks of motility data with respect to tip fluorescence per DF (block bootstrap). N =1000 randomized datasets. E. Frequency histogram displaying distribution of TCS CCFs for randomized subcluster datasets (grey bars) compared to observed TCS CCF (color dashed line). Frequency of randomized mean peak CCF exceeding observed mean peak CCF: EGFP-MENA 114/1000 (bootstrap p = 0.114); EGFP-EVL 3/1000 (bootstrap p = .003). See also Fig. 3.

**Supplemental Figure 4:**
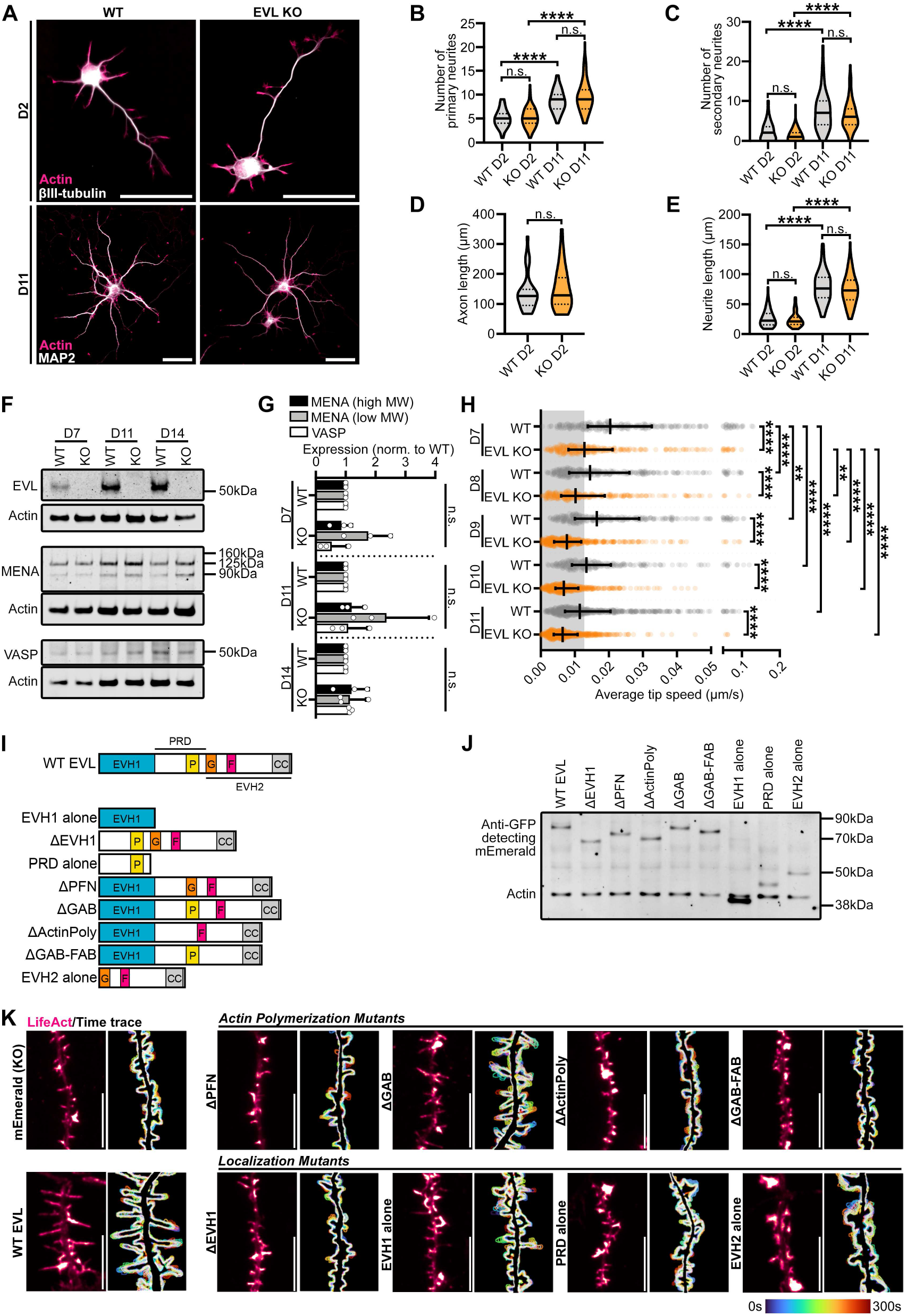
EVL regulates DF morphogenesis and motility through membrane-targeted actin polymerization. A. Immunofluorescence labeling of primary cortical neurons derived from wildtype (WT) or EVL knockout (KO) mice at indicated days *in vitro*. β-III-tubulin labeling shows overall neurite and early axon morphology, while MAP2 specifically labels dendrites. Scale bar = 50µm. B-E Quantification of neurite morphogenesis in cortical neurons derived from WT or EVL KO mice at indicated days *in vitro*. B. Number of primary neurites originating from the soma. C. Number of higher-order branches (secondary or higher). D. Early axon length. The presumptive axon was defined as a primary neurite with a length greater than three times longer than the minor neurites. E. Average length of primary neurites (excluding presumptive axons). Central line = median, dashed lines = interquartile range (IQR). F-G Representative western blot (F) and quantification of (G) protein lysates from cortical neurons derived from WT or EVL KO mice at indicated days *in vitro*, demonstrating complete loss of EVL, and no compensatory upregulation of VASP or MENA high and low molecular weight isoforms. H. Scatterplot of average speed of DF tips, calculated as the average absolute tip displacement between successive timepoints. Grey shaded region indicates average speed less than 0.0128µm/s (non-motile DFs). Median +/- IQR. I. Schematic of protein domains of wildtype EVL and EVL mutants used in this study. EVH1 (blue) = Enabled/VASP homology-1, P (yellow) = profilin binding region, G (orange) = G-actin binding region, F (red) = F-actin binding region, CC (grey) = coiled-coil domain, PRD = proline-rich region, EVH2 = Enabled/VASP homology-2. J. Western blot of protein lysates from D11 cortical neurons derived from EVL KO mice expressing mEmerald-EVL or various mutants of EVL as indicated. K. Segment of dendrite from live EVL KO cortical neurons at D11 expressing mRuby2-LifeAct and mEmerald, wildtype, or mutant mEmerald-EVL as indicated. Left: LifeAct presented with an intensity-coded LUT. Right: maximum intensity projection of temporally color-coded binary mask outline (5s interval, 5min duration). Scale bar = 10µm. Statistical comparisons were made as follows: (B,C,E) Kruskal-Wallis test and corrected for multiple comparisons. n = 77-99, N = 3 biological replicates. (D) Mann-Whitney test. n = 32-35, H: n = 248-397, N = 3 biological replicates. (G) made by unpaired t-test. N = 3 biological replicates. (H) Kruskal-Wallis test and corrected for multiple comparisons. n = 248-397, N = 3-5 biological replicates. * P < 0.05, ** P < 0.01, *** P < 0.001, **** P < 0.0001, n.s. is not significant. See also Fig. 4.

**Supplementary Figure 5:**
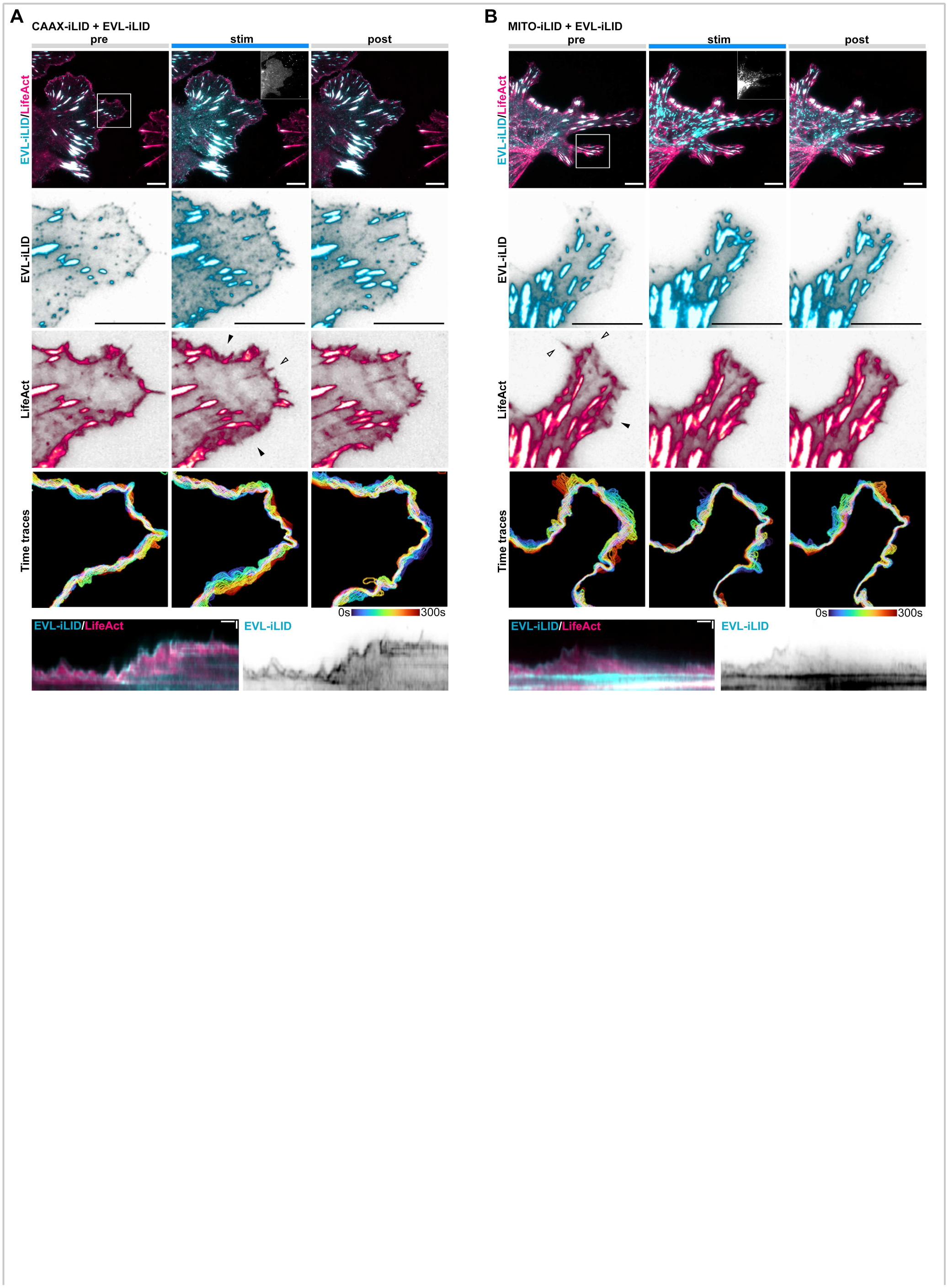
EVL tip localization is necessary and sufficient for DF motility. A-B Glial cells expressing iRFP670-LifeAct, EVL-iLID, and either CAAX-iLID (A) or MITO-iLID (B). Cells were photostimulated with 488nm light for 5min, and imaged for 5min before, during, and after photostimulation. Row 1: localization of LifeAct and EVL-iLID during indicated phases of photostimulation. Inset: mVenus-CAAX-iLID or mVenus-MITO-iLID localization. Scale bars = 10µm. Rows 2-3: localization of EVL-iLID and LifeAct in magnified region of lamellipodia indicated in Row 1. Individual channels displayed with black subtraction for ease of morphological comparison. Arrowheads indicate regions which exhibited altered dynamics following photostimulation (lamellipodia by empty arrowheads, filopodia by filled arrowheads). Row 4: maximum intensity projection of temporally color-coded binary mask outline. Row 5: kymograph of cell’s edge. 15min duration, 5s interval, vertical scale bar = 1µm, horizontal scale bar = 1min, dashed line indicates dendrite. See also Fig. 5.

**Supplementary Figure 6:**
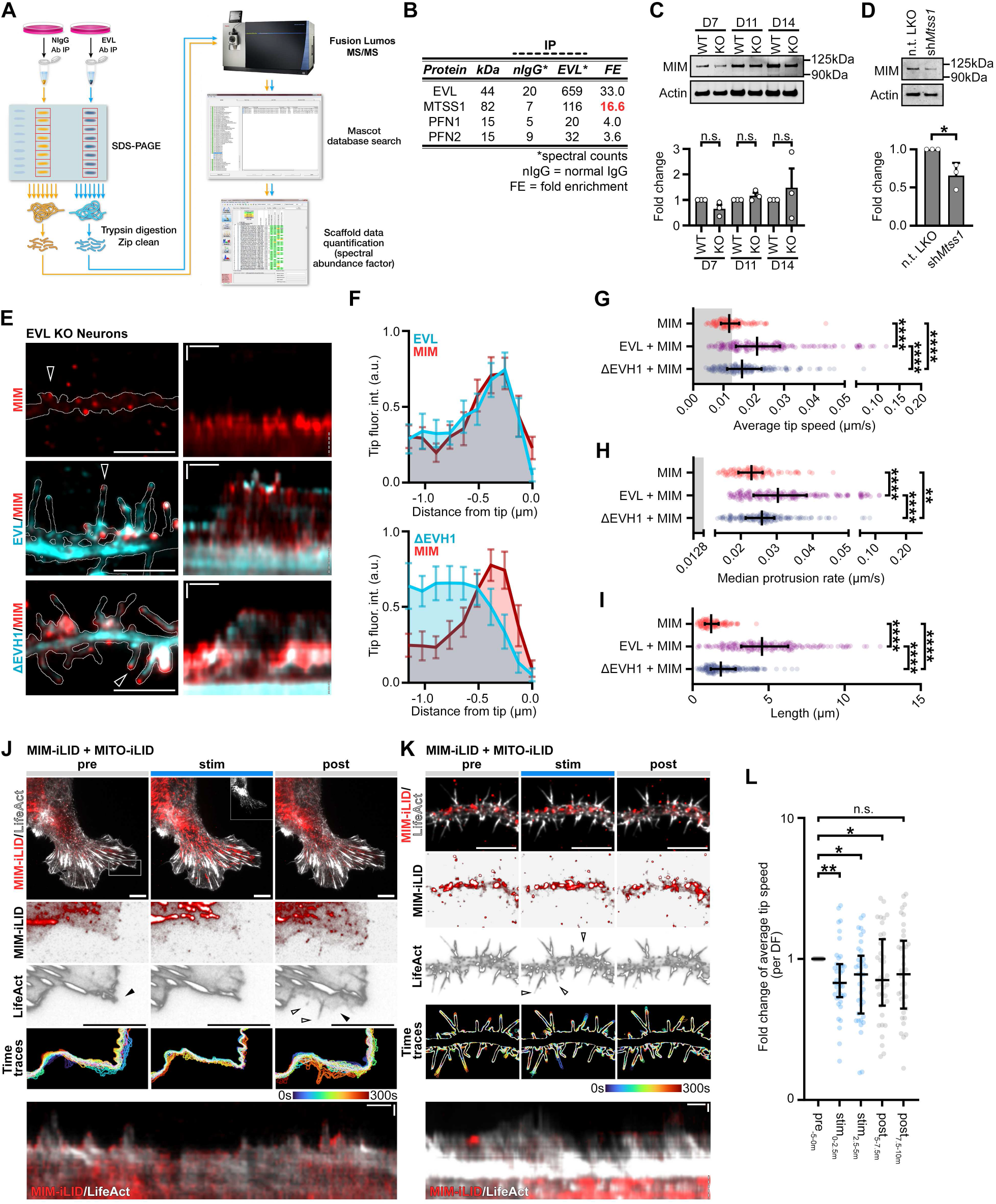
MIM/MTSS1 cooperates with EVL to promote DF initiation and motility. A. Schematic of affinity purification-mass spectrometry (AP-MS) workflow to identify putative partners of endogenous EVL. B. Table of select actin-regulating proteins identified by AP-MS. C. Representative western blot and quantification of protein lysates from cortical neurons derived from WT or EVL KO mice at indicated days *in vitro*, and probed with an antibody targeting MIM. D. Representative western blot of protein lysates from primary neurons at D11, transduced on D7 with indicated pLKO-shRNA-TurboRFP lentiviral particles targeting *Mtss1* or non-targeting (n.t.) control. Fold change quantification of knockdown compared to non-targeting shRNA control. E. Segment of dendrite from live EVL KO cortical neurons at D11 expressing MIM-mRuby2, or MIM together with mEmerald-tagged wildtype EVL or ΔEVH1 as indicated (left). Right: kymograph of DF position indicated by arrowhead in left column. 5s interval, 5min duration, dendrite segment: scale bar = 10µm. Kymograph: Horizontal scale bar = 1min, vertical scale bar = 1µm. F. Distribution of MIM and EVL or ΔEVH1 localization along the distal length of DF. Mean +/- 95% confidence interval. G. Scatterplot of average speed of KO DF tips for indicated conditions, calculated as the average absolute tip displacement between successive timepoints. Grey shaded region indicates average speed less than 0.0128µm/s (non-motile DFs). Median +/- interquartile range (IQR). H. Scatterplot of median protrusion rates of KO DFs for indicated conditions (the median of values when instantaneous change in length was greater than +0.0128µm/s (motile, protruding)). Median +/- IQR. I. Scatterplot of average length of KO DFs reached during the duration of imaging for indicated conditions. Median +/- IQR. J,K Glial cells (J) and cortical neurons (K) derived from WT mice expressing iRFP670-LifeAct, MIM-iLID, and MITO-iLID. Cells were photostimulated with 488nm light for 5min, and imaged for 5min before, during, and after photostimulation. Row 1: localization of LifeAct and MIM-iLID during indicated phases of photostimulation. Inset: mVenus-MITO-iLID localization (J only). Rows 2-3: localization of MIM-iLID and LifeAct in magnified region of lamellipodia indicated in Row 1 (J) or the full dendrite segment (K). Individual channels displayed with black subtraction for ease of morphological comparison. Arrowheads indicate regions which exhibited altered dynamics following photostimulation (lamellipodia= filled arrowheads, filopodia = empty arrowheads). Row 4: maximum intensity projection of temporally color-coded binary mask outline. Row 5: kymograph of cell’s edge (J) or representative DF position (K). Scale bar = 10µm. 5s interval, 15min duration. L. Scatterplot of fold change of average tip speed, calculated as the average absolute tip displacement between successive timepoints relative to average tip speed pre-photostimulation, for 2.5min bins during or post-photostimulation, for WT neurons expressing MIM-iLID with MITO-ILID. Median +/- IQR. Statistical comparisons were made as follows: (C,D) t-test. N = 3 biological replicates. (G-I) Kruskal-Wallis test and corrected for multiple comparisons. n = 101-195, N = 3 biological replicates. (L) were paired and followed individual DFs during phases of stimulation, using Friedman’s Test. n = 39, N = 2 biological replicates. * P < 0.05, ** P < 0.01, *** P < 0.001, **** P < 0.0001, n.s. is not significant. See also Fig. 6.

## Supplementary Video Legends

**Supplementary Video 1:** Live primary mouse cortical neurons at day *in vitro* 11 (D11) expressing EGFP-LifeAct and pLKO-shRNA-TurboRFP targeting *Enah, Evl*, or non-targeting (n.t.) as indicated. Left: segment of dendrite with overlay of brightfield and intensity-coded LUT of EGFP-LifeAct. Right: EGFP-LifeAct alone. Scale bar = 10µm. 5s interval, 5min duration.

**Supplementary Video 2:** Live primary mouse cortical neurons at day *in vitro* 11 (D11) expressing mRuby2-LifeAct and EGFP (top row), EGFP-MENA (middle row), or EGFP-EVL (bottom row). Left: segment of dendrite with merge of EGFP (cyan) and LifeAct (magenta). Right: rainbow intensity-coded LUT of EGFP with outline of LifeAct-positive borders. Red = highest intensity, purple = lowest intensity. Scale bar = 10µm. 5s interval, 5min duration.

**Supplementary Video 3:** Live primary cortical neurons derived from wildtype (WT) or EVL knockout (KO) mice at indicated days *in vitro* expressing mRuby2-LifeAct. Left: segment of dendrite with overlay of brightfield and intensity-coded LUT of mRuby2-LifeAct. Right: mRuby2-LifeAct alone. Scale bar = 10µm. 5s interval, 5min duration.

**Supplementary Video 4:** Live EVL KO cortical neurons at day *in vitro* 11 (D11) expressing mRuby2-LifeAct and mEmerald alone, or mEmerald-tagged wildtype EVL or mutants of EVL as indicated. Phase 1: segment of dendrite with merge of mEmerald (cyan) and LifeAct (magenta). Phase 2: intensity-coded LUT of mRuby2-LifeAct. Phase 3: rainbow intensity-coded LUT of mEmerald with outline of LifeAct-positive borders. Red = highest intensity, purple = lowest intensity. Scale bar = 10µm. 5s interval, 5min duration.

**Supplementary Video 5:** Live primary EVL KO mouse cortical neurons at day *in vitro* 11 (D11) expressing iRFP670-LifeAct and indicated iLID constructs. Neurons were photostimulated with 488nm laser light for 5min, and imaged for 5min before, during, and after photostimulation (stimulation period indicated by blue circle). Row 1: segment of dendrite with merge of indicated EVL-iLID constructs (cyan) and LifeAct (magenta). Row 2: iLID localization constructs CAAX-iLID or MITO-iLID (images acquired only during stimulation). Row 3: rainbow intensity-coded LUT of indicated EVL-iLID constructs with outline of LifeAct-positive borders. Red = highest intensity, purple = lowest intensity. Row 4: intensity-coded LUT of iRFP670-LifeAct. Scale bar = 10µm. 5s interval, 15min duration.

**Supplementary Video 6:** Live primary WT mouse cortical neurons at day *in vitro* 11 (D11) expressing mRuby2-LifeAct, together with mEmerald-EVL and/or MIM-iRFP670 as indicated. Left: segment of dendrite with overlay of brightfield and intensity-coded LUT of mRuby2-LifeAct. Right, bottom: rainbow intensity-coded LUT of indicated constructs with outline of LifeAct-positive borders. Red = highest intensity, purple = lowest intensity. Right (MIM+EVL only): merge of mEmerald-EVL (cyan) and MIM-iRFP670 (red). Scale bar = 10µm. 5s interval, 5min duration.

**Supplementary Video 7:** Live primary WT mouse cortical neurons at day *in vitro* 11 (D11) expressing EVL-iLID and MIM-iRFP670, and CAAX-iLID or MITO-iLID as indicated. Neurons were photostimulated with 488nm laser light for 5min, and imaged for 5min before, during, and after photostimulation (stimulation period indicated by blue circle in row 2). Row 1: segment of dendrite with merge of EVL-iLID (cyan), MIM-iRFP670 (red), and brightfield to better display DF dynamics. Row 2: merge of EVL-iLID (cyan), MIM-iRFP670 (red) only. Row 3 and 4: rainbow intensity-coded LUT of EVL-iLID (row 3) or MIM-iRFP670 (row 4). Red = highest intensity, purple = lowest intensity. Scale bar = 10µm. 5s interval, 15min duration.

**Supplementary Video 8:** Segments of dendrite (left, middle) or dendrite growth cone (right) from live primary cortical neurons derived from wildtype (WT) or EVL KO mice at day *in vitro* 11 (D11) expressing mRuby2-LifeAct. Cultures were treated with 0.1% DMSO, 100µM CK-666, or 10nM SMIFH2 at 5 minutes, and imaged for 40 minutes following treatment (treatment period indicated by yellow square). Scale bar = 10µm. 15s interval, 45min duration.

## KEY RESOURCES TABLE

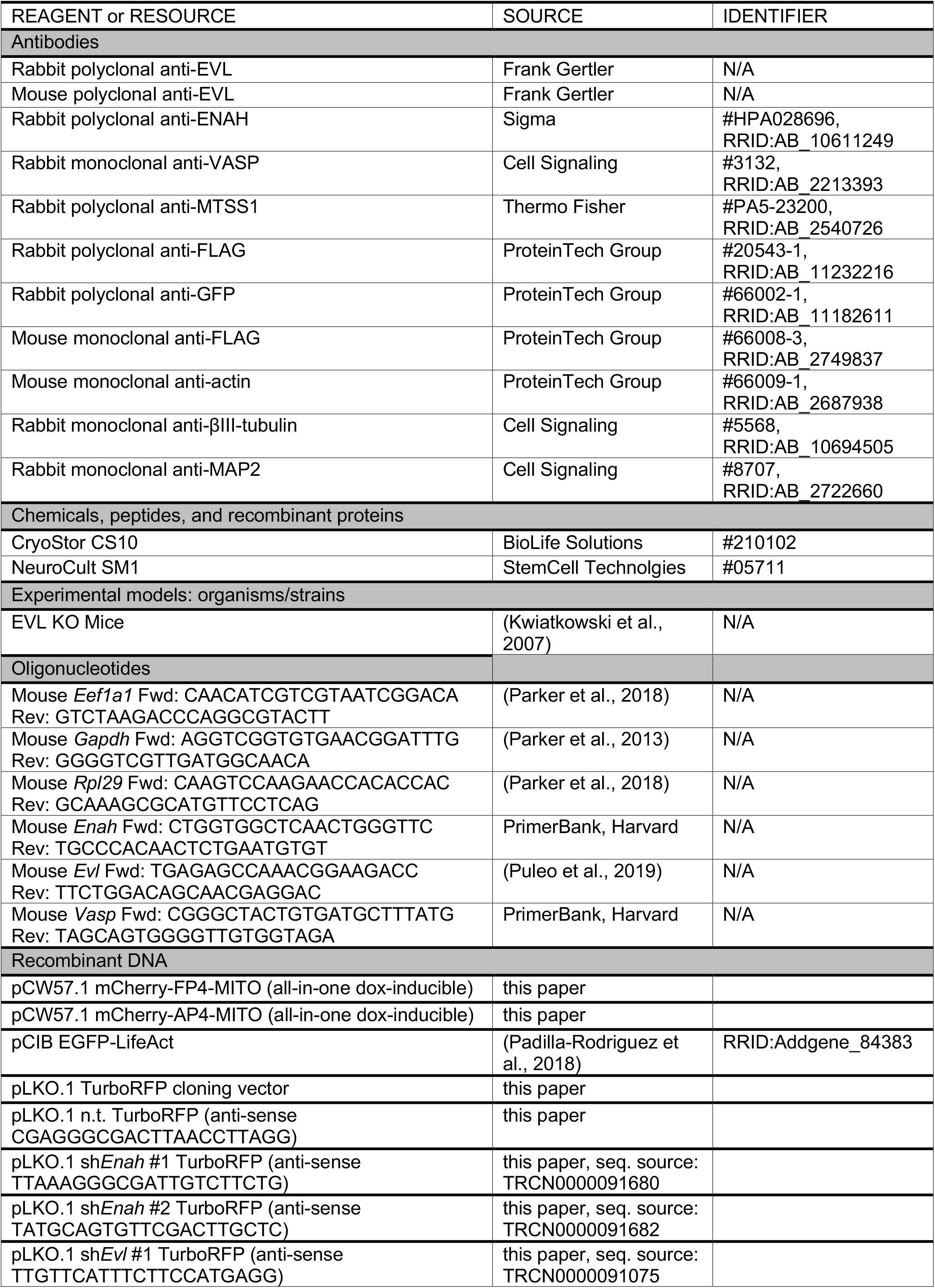
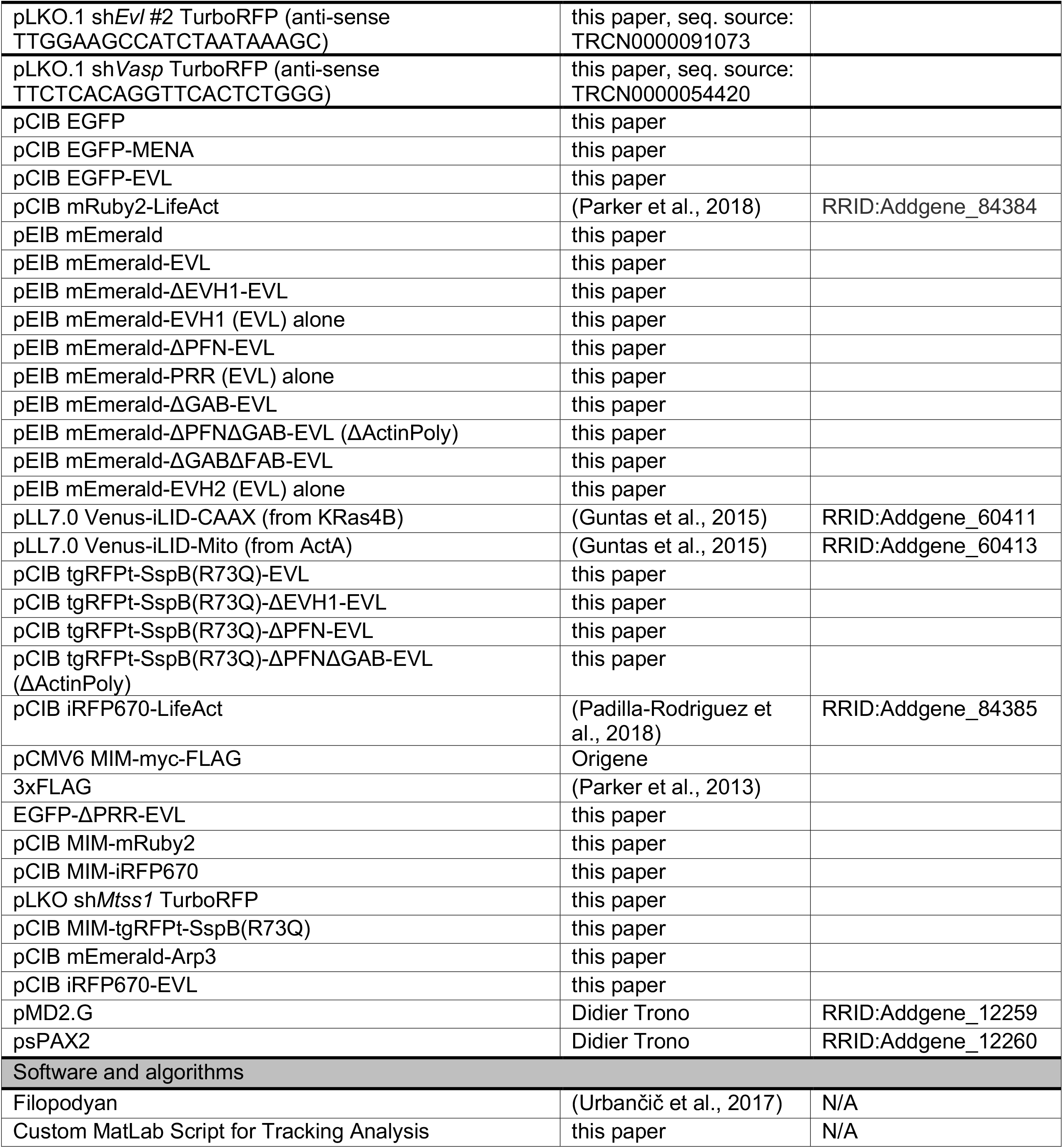

## Notes

### Competing Interest Statement

The authors have declared no competing interest.

